# eLemur: A cellular-resolution 3D atlas of the mouse lemur brain

**DOI:** 10.1101/2024.07.05.601641

**Authors:** Hyungju Jeon, Jiwon Kim, Jayoung Kim, Yoon Kyoung Choi, Chun Lum Andy Ho, Fabien Pifferi, Daniel Huber, Linqing Feng, Jinhyun Kim

**Affiliations:** Brain Science Institute, Korea Institute of Science and Technology (KIST), Seoul, South Korea; Division of Bio-Medical Science & Technology, KIST-School, University of Science and Technology, Seoul, South Korea; Department of Computer Science and Engineering, Korea University, Seoul, South Korea; University of Geneva, Department of Basic Neurosciences, Geneva, Switzerland; Musée National d’Histoire Naturelle, Adaptive Mechanisms and Evolution, UMR7179—CNRS, Paris, France; KIST-SKKU Brain Research Center, SKKU Institute for Convergence, Sungkyunkwan University, Suwon, South Korea

**Keywords:** Grey mouse lemur (Microcebus murinus), 3D brain atlas, cellular resolution, website resources

## Abstract

The gray mouse lemur (*Microcebus murinus*), one of the smallest living primates, emerges as a promising model organism for neuroscience research. This is due to its genetic similarity to humans, its evolutionary position between rodents and humans, and its primate-like features encapsulated within a rodent-sized brain. Despite its potential, the absence of a comprehensive reference brain atlas impedes the progress of research endeavors in this species, particularly at the microscopic level. Existing references have largely been confined to the macroscopic scale, lacking detailed anatomical information. Here, we present *eLemur*, a new resource, comprising a repository of high-resolution brain-wide images immunostained with multiple cell type and structural markers, elucidating the cyto- and chemoarchitecture of the mouse lemur brain. Additionally, it encompasses a segmented two-dimensional (2D) reference and 3D anatomical brain atlas delineated into cortical, subcortical, and other vital regions. Furthermore, eLemur includes a comprehensive 3D cell atlas, providing densities and spatial distributions of non-neuronal and neuronal cells across the mouse lemur brain. Accessible via a web-based viewer (https://eeum-brain.com/#/lemurdatasets), the eLemur resource streamlines data sharing and integration, fostering the exploration of new hypotheses and experimental designs using the mouse lemur as a model organism. Moreover, in conjunction with the growing 3D datasets for rodents, non-human primates, and humans, our eLemur 3D digital framework enhances the potential for comparative analysis and translation research, facilitating the integration of extensive rodent study data into human studies.

**Significance Statement:** The gray mouse lemur (*Microcebus murinus*) represents a promising model for neuroscience research, offering insights into brain structure and function due to its genetic and evolutionary proximity to humans and rodents. Our development of *eLemur*, a comprehensive 3D digital brain atlas, fills critical gaps in microscopic data availability, enabling nuanced investigations at cellular resolutions. This resource not only enhances our understanding of the mouse lemur brain but also catalyzes broader neuroscience endeavors. By facilitating data sharing and integration, eLemur empowers researchers to explore novel hypotheses and experimental designs. Moreover, its compatibility with existing neuroanatomy frameworks and the growing repository of 3D datasets positions eLemur as a pivotal tool for advancing basic and translational neuroscience studies.

## Introduction

Model organisms have been pivotal in propelling advancements in biology, especially in the realm of modern neuroscience research^1–8^. The laboratory mouse, in particular, has led to significant breakthroughs in genetics and cutting-edge technologies (from gene editing to automated behavioral analyses), shaping our understanding of the brain^9–14^. However, despite these strides, challenges persist in translating rodent-based findings to primate biology, behavior, and human-relevant diseases. Neuroscience research often is conducted under the assumption that findings generalize across species, but many brain areas, including the neocortex, have species-specific functional organizations. Recognizing the need for diverse animal models and a comparative approach, there is growing interest in expanding the portfolio of model species for next-generation neuroscience research^5,7,15–19^.

Among emerging candidates, the mouse lemur (*Microcebus murinus*) has drawn attention as a promising novel non-human primate model system^18,20–25^. Mouse lemurs are suggested as a practical and cost-efficient model for neuroscience research due to their unique attributes, such as its small rodent-like body and brain size, potential for rapid colony growth, and genetic proximity to humans. Moreover, the mouse lemur exhibits remarkable adaptability to laboratory settings and holds the potential for leveraging transferable tools developed for mice, positioning it as an advantageous primate model system that bridges the gap between primate and rodent studies^18^. Studies employing this species increase, exploring comparable information and transferable experimental paradigms between the mouse lemur, primates, and rodents^23,26–29^. These endeavors support the potential of the mouse lemur for elucidating fundamental aspects of neurobiology, ultimately facilitating the translation of extensive rodent studies into organizational principles governing further complex nervous systems.

To establish the mouse lemur as a new model organism, our understanding of its genetic and anatomical features is fundamental. Currently, ongoing genome sequencing projects aim to extensively map the mouse lemur genome^30^, while single-cell resolution transcriptome projects strive to provide comprehensive molecular cell profiles^22^. Additionally, classic Nissl stain-based stereotaxic atlases and MRI-based brain atlases have already been established^27,31,32^. Nevertheless, these atlases, while informative, provide only a coarse view of anatomical structures lacking the spatial molecular and cellular details crucial for functional inference. Identifying brain subregions and specific cell types is essential for elucidating functional diversity, mapping neural circuits, and uncovering the cellular basis of neurological disorders. Ultimately, this knowledge will deepen our comprehension of brain function and dysfunction, guiding targeted therapeutic interventions^33^.

In addressing this gap, we present a digital 2D/3D atlas of the mouse lemur brain with cellular-resolution, incorporating advanced fluorescence immuno-labeling and computational imaging analysis. This digital atlas, named *eLemur*, is publicly accessible through our web-based viewer (https://eeum-brain.com/#/lemurdatasets). eLemur comprises: (1) a repository of high-resolution brain-wide immuno- fluorescence images with multiple cell type and structural markers, revealing the cyto- and chemoarchitecture of the mouse lemur brain; (2) a 2D/3D brain atlas serving as a template, segmented into 54 brain subregions based on a hierarchical ontology; and (3) a 3D cell atlas providing region-by-region densities and positions of non-, neuronal, and parvalbumin expressing cells. Our cellular-resolution 3D atlas of the mouse lemur brain, eLemur, offers a resource for the neuroscience community and will facilitate future research bridging gaps in our understanding of rodents, non-human primates and humans.

## Results

### Mouse lemur brain atlas pipeline and data resources

To construct a high-resolution 3D atlas of the mouse lemur brain, we devised a pipeline comprising several key steps: whole-brain sectioning with block face imaging, brain-wide multiplex immunolabeling and fluorescence imaging with subsequent imaging processing, brain region annotation, 2D image alignment, cell detection analysis leading to 3D atlas generation, and implementation of an interactive web-based visualization platform (**Fig. 1A; see Methods**). First, we explored suitable markers capable of delineating brain structures, axonal projections, and cell types compatible with mouse lemur brain tissue. Following a thorough comparison with mouse brain tissue, we selected a total of nine markers, along with DAPI, exhibiting high specificity in the mouse lemur brain to generate brain-wide fluorescence immunohistochemistry data (**Table S1**). The mouse lemur brains were coronally sectioned in 50 µm thickness, obtaining approximately 360 sections per brain. We conducted multiplex immunofluorescence staining with combinations of markers on serial coronal sections in order to intricately delineate the structural features of brain subregions (see **Fig. S1**; **Methods**). For subsequent registration of immunofluorescence images in the step of 3D atlas generation, the surface of the brain tissue block was captured using a digital camera before each sectioning process (referred to as block face image)^34^. The high-resolution combinatorial multiplex immunofluorescence datasets, with a resolution of 0.65 µm in x-y dimensions, enabled the exploration of the intricate cyto- and chemoarchitecture of the mouse lemur brain. Immunofluorescence signals from the neuron-specific nuclear protein (NeuN), coupled with DAPI counterstaining, provided an overview of each the neuronal and non-neuronal population across different brain regions. Moreover, datasets obtained from parvalbumin (PV), tyrosine hydroxylase (TH), and Forkhead box protein P2 (FOXP2) staining revealed the spatial distribution of diverse cell types across the brain, thereby facilitating the analysis of distinct neural components (**Fig. 1B, C**). Furthermore, TH signals were instrumental in characterizing the basal ganglia (BG), a group of subcortical nuclei implicated in motor control and movement disorders. Additionally, signals derived from vesicular glutamate transporter 2 (VGLUT2) and myelin basic protein (SMI-99) allowed for the differentiation of subregional structures in the cortex, hippocampus, and sensory thalamus (**Fig. 1B**, **Fig. 2**). These selected markers collectively serve as neuroscientific probes, enabling the investigation of distinct neural structures of the mouse lemur brain. Through our initial investigation, we uncovered anatomical homologies shared across primate species, underscoring the mouse lemur’s potential as a bridging model between rodent and primate neuroscience studies (**Fig. 1D; Fig. S2**). Of notable significance is cortical folding and the presence of the caudate nucleus and putamen, separated by a distinct fiber tract, a feature commonly observed in primate brains^35^. While in rodents, these structures often appear fused into a unified striatum, in primates, they serve distinct roles in goal-directed behavior and sensorimotor coordination^36^. Additionally, we observed the globus pallidus interna (GPi) as a distinct structure adjacent to the globus pallidus externa (GPe), a configuration differing from rodents where the entopeduncular nucleus (EPN) is considered a debated homolog of the primate GPi^37,38^. The lateral geniculate nucleus (LGN), also known as the visual thalamus in mouse lemurs, exhibits a distinct six-layered structure, indicating a sophisticated visual information processing system that is common among primates but absent in rodents. Furthermore, in mouse lemurs, as in primates, the primary visual cortex exhibits a distinct division of layer 4 into 4a and 4b (**Fig. 2A, B**). Each of these sublayers receives primary inputs from the LGN, conveying information from the contralateral and ipsilateral eye. This characteristic is unique to primates and is also observed in mouse lemurs, in contrast to rodents, where a single layer 4 receives thalamic inputs^26,39^. Our primary objective was to establish a repository for brain-wide immunofluorescence datasets, aiming to offer fundamental insights into the intricate neural architecture and molecular characteristics of the mouse lemur brain. These findings underscore the mouse lemur’s potential as a promising model organism for understanding neural complexities across primate species. In conjunction with the previous study revealing hundreds of primate-specific gene expressions absent in mice^22^, these brain anatomical findings highlight the mouse lemur’s unique position as an intermediary model between rodents and primates, offering broader translational implications for understanding human brain functions and associated disorders.

**Figure 1.**
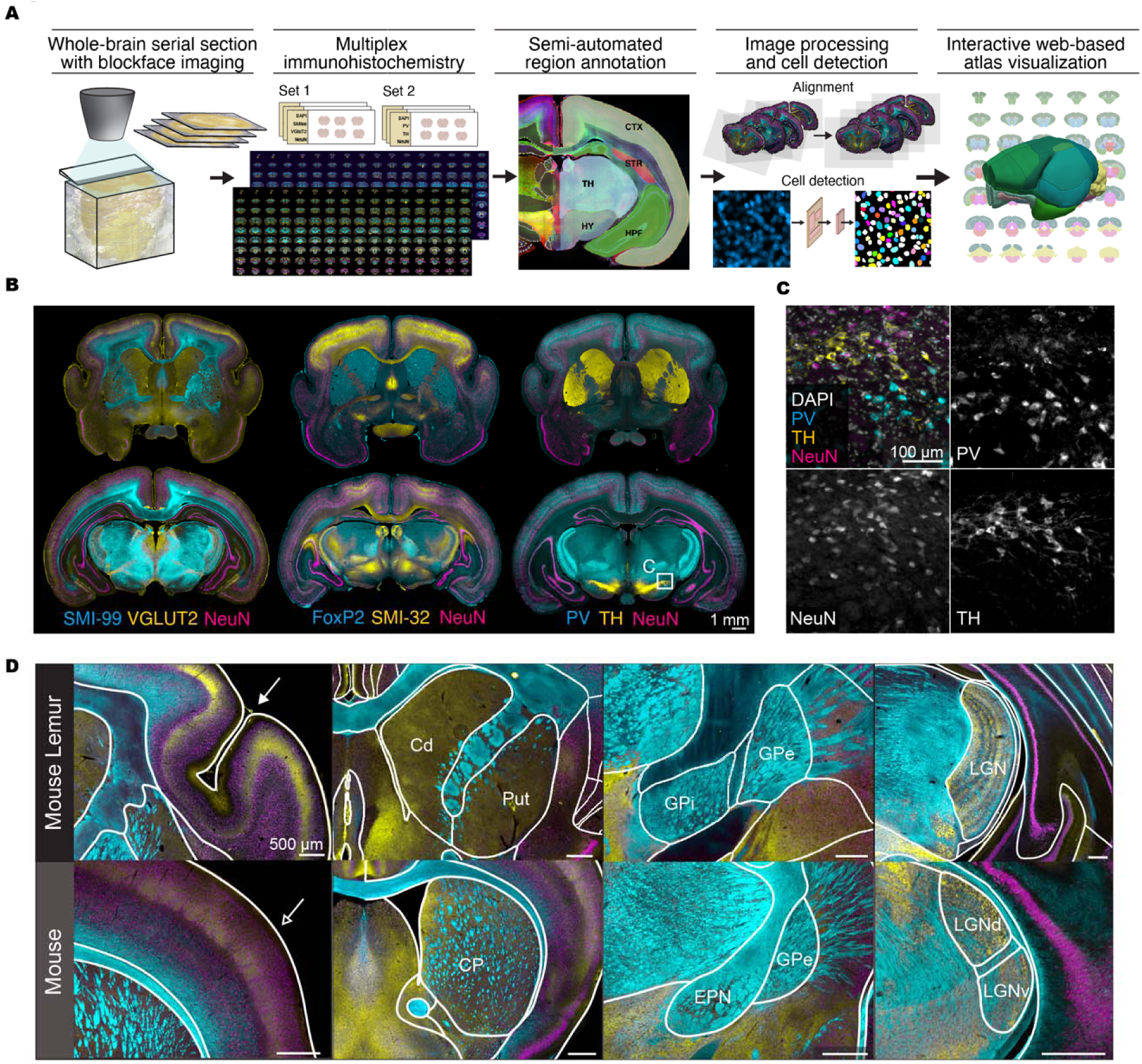
Atlas pipeline and anatomical features of the mouse lemur brain. **(A)** Illustration of the pipeline generating multiplex brain-wide immunofluorescence image datasets and constructing a 2D/3D atlas of the mouse lemur brain. **(B)** Representative immunofluorescence images showcasing multiplex cell type and structural markers, illustrating the cytoarchitecture of the mouse lemur brain. **(C)** High magnification images of the boxed area in (B), displaying NeuN-, PV-, and TH-positive cell types in the substantia nigra. **(D)** Comparison of anatomical features between the mouse lemur and mouse brains. The mouse lemur brain exhibits anatomical homologies shared with other primates, including cortical folding (top left, indicated by an arrowhead), the separated caudate and putamen, as well as the GPi (top middle), and the laminar structure of the LGN (top right). These features are absent in the mouse brain (bottom).

**Figure 2.**
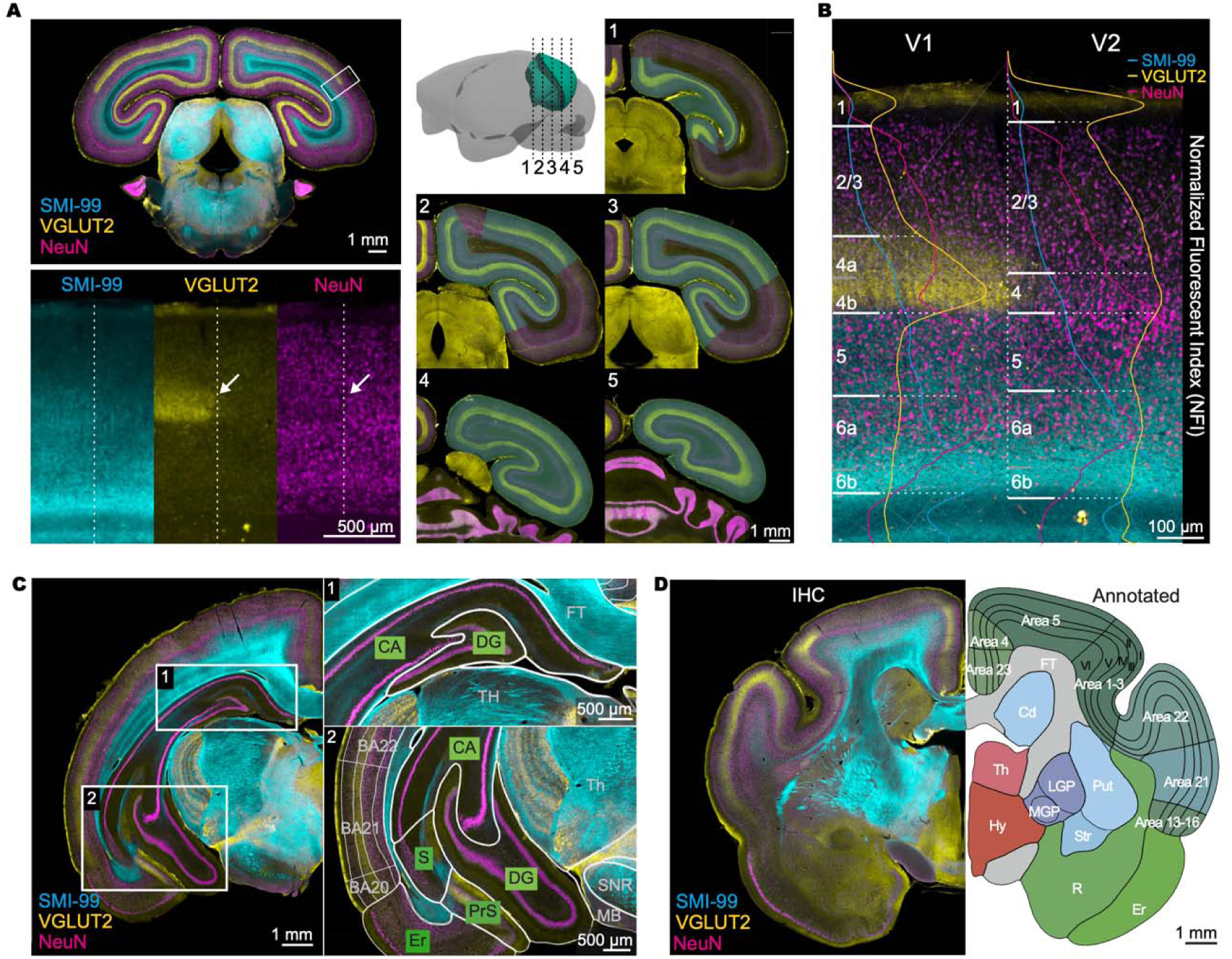
Delineating brain structures and annotating a 2D reference atlas. **(A)** Representative coronal immunofluorescence image with three marker staining: SMI-99, VGLUT2, and NeuN. Enlarged images of the boxed area at the V1-V2 border (left). The boundary between V1-V2, indicated by a dashed line, is defined by the sharp transition of VGLUT2 and NeuN signals (indicated by an arrow). The progression of the V1 area across the longitudinal axis (anterior-posterior) is shown with manual annotations in green (right). **(B)** Cortical layers in V1 and V2 are defined by curated intensity and density of SMI-99, VGLUT2, and NeuN signals. The intensity plot of normalized fluorescence index along a perpendicular cortical column in V1 and V2 for each marker—SMI-99 (cyan), VGLUT2 (yellow), and NeuN (magenta)—is overlaid to demonstrate the laminar structure. **(C)** Hippocampal substructures, including dentate gyrus, subiculum, and entorhinal cortex, are identified and segmented using multiplexed immunosignals. **(D)** Representative 2D coronal section showing full annotations (right) alongside the original immunofluorescence image (left).

### Charting brain structures: Annotating a 2D Reference Atlas

To generate a 2D Reference Atlas of the mouse lemur brain, we annotated a single reference specimen by drawing on its stained images, as traditionally done for the mouse and human^40,41^. The single reference brain was coronally sectioned and alternately divided into two staining sets. Set 1 was stained with NeuN, SMI-99, VGLUT2, and DAPI, while Set 2 was stained with NeuN, PV, TH, and DAPI. The 2D- immunofluorescence images were aligned to corresponding parts in the block face image via 2D rigid multi-scale registration, followed by shading artifact correction (**Fig. S3**). Brain structures were segmented based on their cellular architectures through these multiple markers (**Fig. 2; Movies 1, 2**), employing semi-automated structure annotation with a priority on preserving data in its original image state with minimal computational processing. This approach significantly reduced manual processing time, comprising steps of sparse manual annotations of the 13 major brain structures (neocortex, hippocampal formation, remaining unspecified cortical regions, striatum, pallidum, thalamus, hypothalamus, midbrain, pons, medulla, cerebellum, fiber tracts, and ventricular system) on the right hemisphere of alternate coronal images, training a deep learning-based region segmentation using the manual annotations for automatic annotation of the entire data set, and detailed manual annotation for refinement and finer subregions (**Fig. S4**).

Manual annotations were facilitated by custom software featuring a graphical user interface (GUI) for easy annotation using cubic spline curves defined by control points, which allowed for detailed refinement. For further subregion delineations, we incorporated additional reference information, including a different set of multiplex immunofluorescence signals with brightfield images, the Nissl-based Bons atlas^31^, and an MRI-based atlas^27^ **(Fig. 2B; S5).** This integration of diverse datasets improved fine region boundary identification and enhanced structural delineation **(Fig. 2D)**. Subregion annotations focused on cortical layers, the hippocampus, and the basal ganglia, considering mouse lemurs’ specific attributes and their important functions.

### Cortical areas and layers

Given their important functions, we aimed for a finer level of annotations for cortical areas. The delineation of cortical layers is crucial for gaining insights into the functional specialization of different layers, which contributes to our understanding of neural circuits and their role in complex cognitive processes. Additionally, precise delineation facilitates comparative studies across species, shedding light on evolutionary adaptations and differences in brain function and behavior. The neocortex’s laminar structure, characterized by distinct molecular compositions and connectivity^42–45^, poses challenges in delineating cortical layers based solely on cellular features, such as those indicated by DAPI and NeuN staining. To address this challenge, we combined cellular features from DAPI and NeuN with layer-specific synaptic terminal distribution indicated by VGLUT2, and integrated annotations from the Bons atlas^31^ to accurately annotate cortical layers (**Fig. 2A, B**). Utilizing VGLUT2 signals was useful in identifying sublayers, such as the dense concentrations in layer 4 and blobs of layer 3 in primary visual cortex (V1 or Brodmann area 17) and secondary visual cortex (V2 or Brodmann area 18)^26,46,47^.

Additionally, VGLUT2 signals helped distinguish sublayers 4a and 4b, with a narrow layer of high cell density and dense VGLUT2-labeled axon terminals in layer 4b and less so in 4a (**Fig. 2B**). Furthermore, we incorporated the MRI-based atlas^27^ to delineate the Brodmann areas, which are specific regions of the cerebral cortex in humans and other primates. This process involved initializing the population averaged T2-weighted MRI template and corresponding labels to the block face volume through a series of 3D affine and non-linear transformations, followed by slice-by-slice 2D deformable registration onto immunofluorescent images (**Fig. S5**). The transferred annotations were then manually refined to ensure that the boundaries between regions aligned parallel to the radial direction. Our detailed annotation of cortical layers and delineation of Brodmann areas in the mouse lemur brain enhances our understanding of the intricate organization and functional specialization of the cerebral cortex across species.

### Hippocampal formation

As learning and memory are fundamental components of higher cognitive functions, the hippocampus (HP) has been extensively studied, with each subregion and layer displaying distinctive organization and function^48^. We undertook segmentation of the hippocampal layers to lay anatomical groundwork for future investigations into hippocampal functions of mouse lemurs, which offer a more complex cognitive model (**Fig. 2C**). Similar to cortical annotations, we curated additional reference information and adopted the Bons atlas. We divided the hippocampal formation into five subregions: the Ammon’s horn (CA), subiculum (S), pre/para-subiculum (PrS), dentate gyrus (DG), and entorhinal cortex (Ent). Each hippocampal subregion consists of a distinguished laminar structure, which formed the basis of our segmentation. The dentate gyrus and Ammon’s horn were identified by densely packed layers of cell bodies, discernible from DAPI and NeuN signals. The boundary between the two regions was delineated by VGLUT2 and SMI-99 signals, along with blood vessels. The subiculum, located adjacent to the Ammon’s horn, was characterized by a relatively sparse pyramidal cell layer, distinctively observable through NeuN staining. The pre/parasubiculum and the entorhinal cortex were distinguished by discontinuation in layer structures.

### Basal ganglia

Despite the well-preserved basic anatomy and connectivity of the basal ganglia across species, interspecific variations exist, notably evidenced from rodents to primates^37,49^. Our inter-species comparison, as depicted in **Fig. 1D** and **S2**, reveals that the mouse lemur exhibits nuanced divergence in key component structures within the basal ganglia, reminiscent of those found in humans. To further elucidate these structures across the brain, we conducted comprehensive segmentation of the basal ganglia, targeting seven key regions implicated in movement control: the caudate, putamen, GPe, GPi, subthalamic nucleus (STh), substantia nigra pars reticulata (SNR), and pars compacta (SNC) (**Fig. 1D**). Employing both the multiplex stainings of VGLUT2/SMI-99/NeuN and TH/PV/NeuN sets, we utilized information primarily from PV and TH staining signals to delineate these substructures. The caudate nucleus and putamen, positioned at the anterior striatum, were separated by the internal capsule, with thin bridges of cells connecting the putamen to the caudate head. The GPi and GPe were located at the medioventral part of the medial- and lateral medullary lamina, respectively. The STh, positioned between the internal capsule and cerebral peduncle, was characterized by its distinctive, bright, lens-shaped nucleus across NeuN, PV, and SMI-32 staining. The pars compacta, located posterior to the STh, could be clearly distinguished from its neighboring regions through TH staining. Meanwhile, the pars reticulata, situated between the pars compacta and large fiber bundles (e.g., cerebral peduncle at the anterior and medial longitudinal fasciculus at the posterior), could be readily identified.

### Ventricular Structures and Fiber Tracts

Utilizing immunofluorescence signals from SMI-99 and NeuN, complemented by brightfield contrast images and annotations from the Bons atlas, we delineated major fiber tracts including white matter structures including the corpus callosum, internal capsule, and cerebral peduncle, were delineated (**Fig. S4A**). These structures, distinguished by darker contrast in the brightfield channel and sparse NeuN signals, were further refined to include fine fiber tracts, such as the mammillothalamic tract within the thalamus. Additionally, within the ventricular system, the lateral, third, and fourth ventricles were annotated based on their distinguishable appearance as gray or empty areas, with ependymal cell-lined walls clearly visualized via DAPI staining, serving as a primary guide for delineation verification and modification (**Fig. S4B**).

### Hierarchical ontology and annotation

Following thorough annotation of the mouse lemur brain, our next step was to optimize the ontology of its brain regions. Our goal was to enhance legibility and ensure compatibility with existing digital frameworks for other species, such as the Allen Reference Atlas (ARA)^50^. We established consistent nomenclature by translating 239 structures from Latin names used in the Bons atlas to English and reconciling inconsistent nomenclatures and abbreviations (see **Table S2**). Our primary references included NeuroNames^51^, a comprehensive neuroanatomical database, and the ARA of the mouse brain when standard names or acronyms were not available in NeuroNames.

To systematically define the relationships between brain regions, we organized a hierarchical ontology based on Brain maps 4.0^52^ and the Allen mouse brain ontology. We structured the mouse lemur brain ontology into a hierarchical tree with four levels. At the root level, divisions were made into grey matter (including cell groups and regions), white matter (fiber tracts), and the ventricular system. The grey matter was further categorized into cerebrum, brain stem, and cerebellum at level 1, which were then subdivided into three hierarchical layers across levels 2-4 (**Table S3**). We defined 13 major brain structures for annotation and further detailed subdivisions of key regions such as the cortical areas, hippocampal formation, and basal ganglia at level 4. To ensure intuitive comparison and compatibility with the widely used ARA, we adopted the color coding of the Allen Institute’s colormap throughout our analysis. A complete hierarchical tree of 54 regions annotated in the reference atlas with unique colors can be found in **Table S3**.

### 3D Reference Atlas of the mouse lemur

The demand for digital 3D atlases is increasing due to their significant advantages over traditional 2D slice-based atlases. These 3D models serve as comprehensive anatomical frameworks, facilitating the integration of findings from various modalities. Deriving precise 3D structures solely from a sequence of 2D atlas images is highly challenging despite provided stereotaxic coordinates, and is prone to misalignment of experimental data due to variations in brain cutting angles. Thus, developing a precise 3D atlas is crucial for understanding the complex anatomy and organization of the mouse lemur brain structures. The eLemur 3D digital atlas address these challenges by providing a common reference model that integrates diverse experimental data, and enables comparative analysis with 3D atlases of other species.

Our 2D to 3D translation process began by registering histological images to corresponding block face images to correct tissue distortions and damages incurred during the sectioning and staining process (**Fig. 3; S3**). Tissue areas from the block face images were semi-automatically labeled, with the correspondence between tissue segments manually determined. Each tissue part in the corrected histology image was aligned with the corresponding tissue parts in the block face image through multi-scale 2D rigid registration. The aligned reference volume provided sufficient spatial information along the z-direction (anterior-posterior) for 3D segmentation and annotation of brain structures. Subsequently, 2D annotations from coronal immunofluorescent images were converted into a 3D format using a combination of automated and manual pipelines (**Fig. 3A**). Transformation parameters, including deformation fields from non-linear registration of coronal images to block face images, were applied to 2D to provide an initial label volume with enhanced smoothness. These annotations were stacked to create labeled volumes, manually corrected for labeling errors, and neighboring slices’ labels were interpolated to yield isotropic volumes (see **Methods**). A 3D mesh was generated from these isotropic label volumes using the marching cubes algorithm^53,54^. Despite careful manual refinement with alignment and interpolation, the generated 3D mesh often exhibited irregularities along the z-axis. To enhance smoothness, Laplacian filters were iteratively applied to each region’s 3D mesh. While this improved visual smoothness, it occasionally led to mesh expansion and overlaps between neighboring regions. In such cases, overlapping regions were identified, and voxels in the overlapping areas were manually unassigned from one of the regions. Each region was iteratively expanded until boundaries met, and this process of smoothing and correcting was repeated until there were no overlapping regions remaining. The eLemur 3D digital atlas offers enhanced spatial comprehension of the intricate anatomy of the mouse lemur brain, thereby advancing our understanding of its structure and function (**Fig. 3B**).

**Figure 3.**
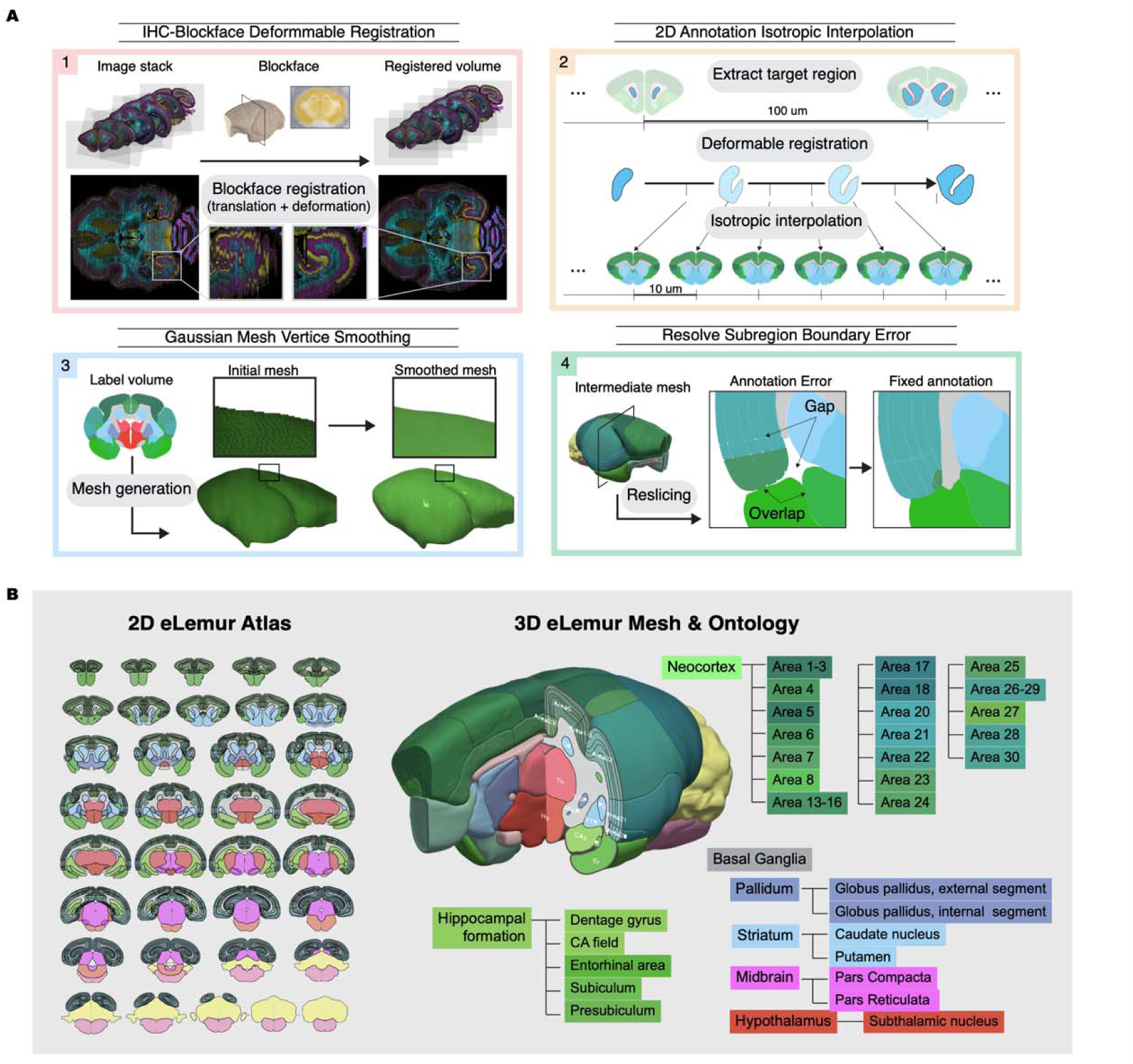
2D/3D reference atlas of the mouse lemur brain. **(A)** Diagram of the process for transforming from 2D to 3D, including block face registration, isotropic interpolation of 2D annotations, and generation of a 3D mesh of brain regions. **(B)** Representative serial annotated 2D atlas (left) and the 3D eLemur mesh with the hierarchical ontology of annotated subregion structures (right). Regions are color-coded based on the color scheme of corresponding structures from the Allen Mouse Brain Atlas.

### 3D brain cell atlas of the mouse lemur

A 3D cell atlas characterizing the distribution of cells across brain regions provides a comprehensive view of the intricate cellular architecture of the brain. Such detailed mapping is essential for deciphering both the functional and structural organization of different brain regions and serves as a fundamental reference for conducting comparative studies across different species. Utilizing digitized datasets, we conducted analysis to quantitatively map region-by-region cellular compositions of the mouse lemur brain, aiming to count total, neuronal, and parvalbumin-positive cells using DAPI, NeuN, and PV-labeled cellular signals, respectively (**Fig. 4A; S6-8; Table S4**). Initially, we developed an automated whole-brain cell detection algorithm with deep learning, employing semi-supervised learning to bootstrap partially annotated data to fully annotated data. After proofreading, the refined fully annotated data trained a Mask R-CNN model with a ResNeXt-101-32x8d^55^ backbone for detecting cells in the entire brain. Validation of our automatic detection results against manually annotated image patches (166.4x166.4 μm) revealed an average precision of 83.9% and a recall rate of 93.8% for DAPI and NeuN detection (**Fig. S7**), while for PV detection, the precision was 81.2% and the recall rate was 96.6% (**Fig. S8**). Furthermore, we benchmarked our model against Cellpose 2.0^56^, where our method exhibited superior performance, possibly due to its reliance on larger training datasets for optimal performance. Since the automatic cell detections were conducted on 50 μm thick 2D images acquired through widefield microscopy, calibration was necessary to estimate total cell counts more accurately in 3D. By imaging subsets of small patches in cortical areas using 3D confocal microscopy and manually counting cells in volumetric images, we found consistent ratios between the number of detected cells in 3D and 2D images across regions (1.730 ± 0.024, SEM for DAPI and 1.864 ± 0.029 for NeuN) (**Fig. 4B, C**). Our automated 3D anatomical template generation pipeline facilitated accurate mapping of detected cell coordinates within 2D images onto 3D reference space.

**Figure 4.**
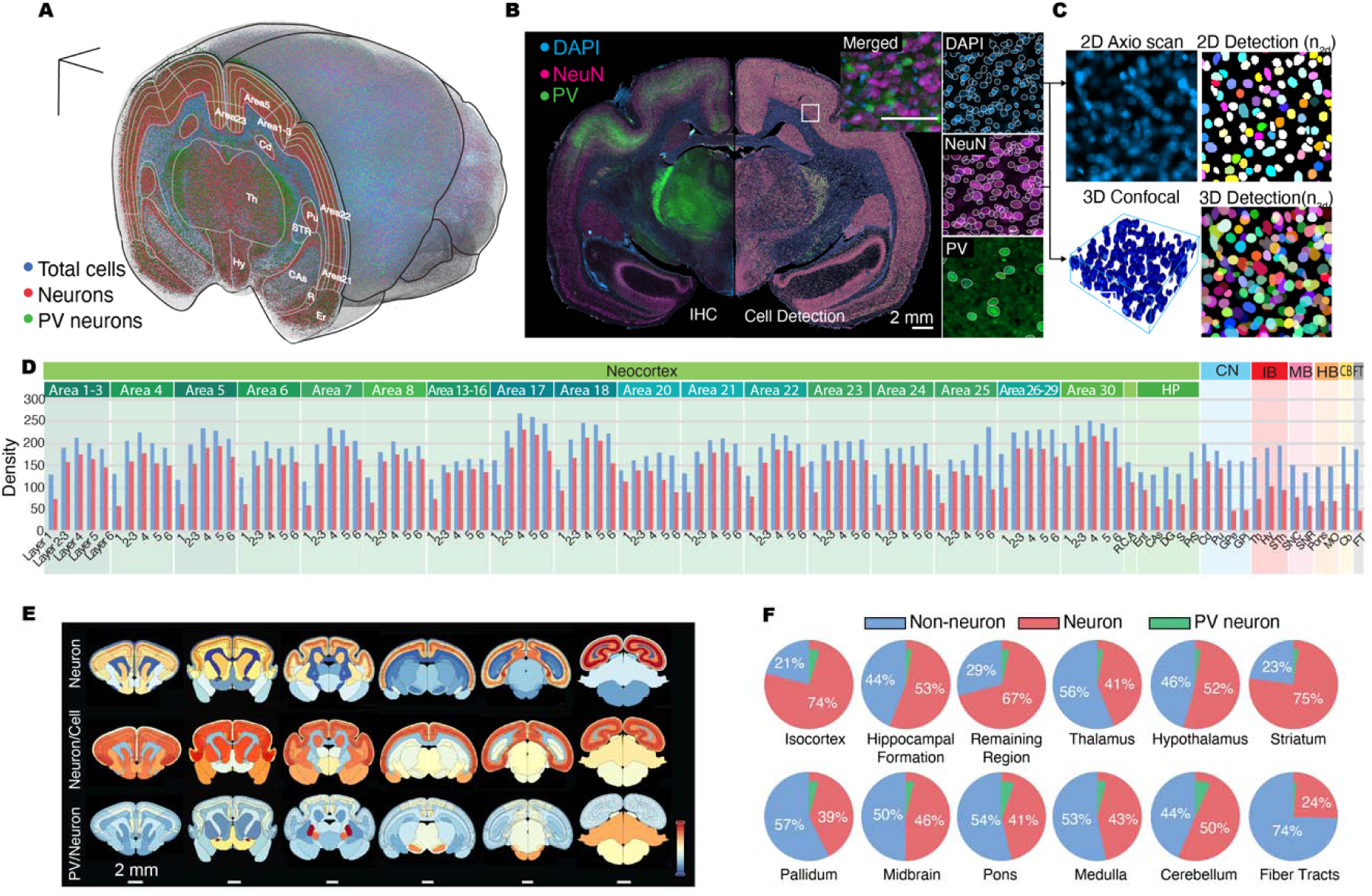
3D brain cell atlas of the mouse lemur brain. **(A)** Cell atlas displaying the topographical distributions of total cells (blue), neurons (magenta), and PV neurons (green) in 3D, labeled by DAPI, anti-NeuN, and anti-PV, respectively. **(B)** Cell detection of NeuN, PV, and DAPI-labeled cells (right) alongside the original immunofluorescence image (left). **(C)** 2D-imaging based cell detection calibrated in 3D by comparing cell count ratios between 3D volumetric confocal images and 2D images. **(D)** Cell composition profiles of non-neuronal (blue) and neuronal (red) cells across brain-wide subregions, including individual cortical layers. Parameter unit: number per 100 μm^3^. **(E)** Heat map of neuronal density (max value: 231.6272; min value 33.6840), neuron/total cell ratio (max value: 1; min value: 0), and PV neuron/neuron (max value: 0.17; min value: 0) ratio across the brain. **(F)** Cell compositions of non-neuronal, neuronal, and PV neurons in twelve major brain structures. The portion of PV neurons is charted in relation to the total number of neurons in each structure.

Consequently, we created a 3D cell atlas of the mouse lemur brain, integrating brain-wide cell estimates of composition and distribution into our 3D atlas database (**Fig. 4D-F**). Detailed 3D-calibrated numbers of the total cells, neurons, PV neurons, and their density of each brain structure can be found in **Table S4** with those of mice^50^.

### Development of a web-based explorer of the mouse lemur

We have developed a comprehensive website for eLemur (**Fig. 5; Movie 3**, https://eeum-brain.com/#/lemurdatasets), offering an immersive and detailed web-based platform that facilitates accessibility, collaboration, data sharing, and visualization for exploring the mouse lemur brain. This interactive platform utilizes the Neuroglancer^57^ engine to enable intricate navigation through high-resolution, multi-terabyte datasets. The design of the eLemur interface aims to provide users with an unrestricted exploratory experience, effectively obviating the need for cumbersome data downloads. The main interface of the portal features a list of curated immunohistochemistry (IHC) datasets, with filtering options available for marker selection (**Fig. 5A**). Each dataset is displayed via a whole-brain gallery view for macroscopic orientation, alongside a full-resolution slice view that permits detailed annotation of brain regions over high-resolution images (**Fig. 5B**). The brain region selection panel facilitates precise navigation and examination of specific areas of interest. Customization of the visualization layout, image channels, and various settings is facilitated through user-operable panels, ensuring a tailored presentation of data. The reference atlas is conveyed through a 2D coronal atlas gallery with accurate physical dimensions (**Fig. 5C**), a cell atlas with regional information of cellular composition **(Fig. 5D)** and a 3D page for interactive mesh visualizations, providing a spatial understanding of structural relationships (**Fig. 5E**).

**Figure 5.**
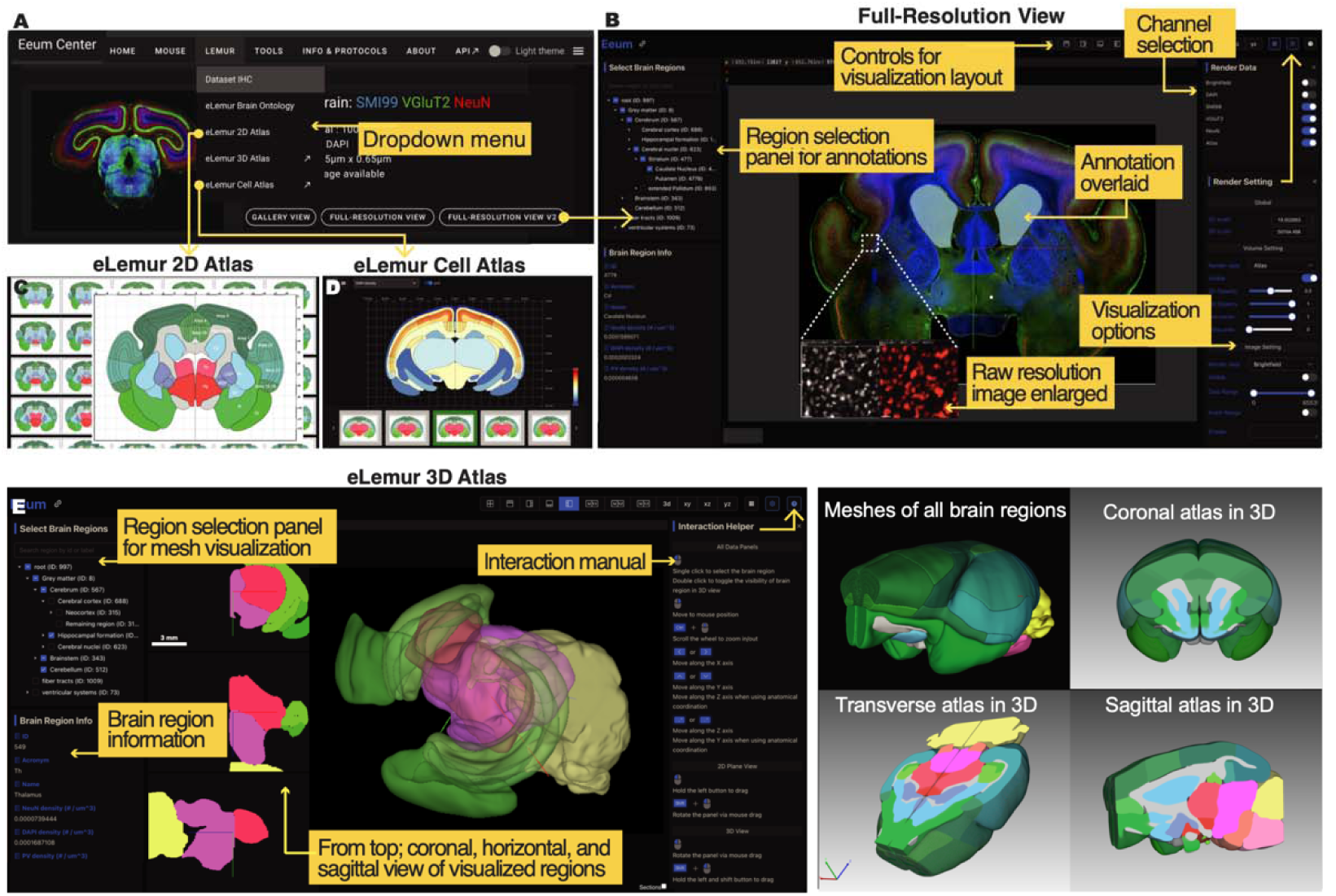
eLemur web-based explorer. **(A)** The dropdown menu from the eLemur panel allows navigation to various datasets. The Dataset IHC panel leads to the repository of immunofluorescence image datasets, featuring tools for antibody-specific filtering and selections, along with whole-brain gallery views for macroscopic orientation. **(B)** High-resolution slice views for detailed examination. Illustrated functionalities include annotation overlays, channel selection, and a suite of visualization settings for customized data representation. **(C)** The eLemur 2D Atlas tab presents a gallery of 2D whole-brain coronal reference images. **(D)** The eLemur Cell Atlas tab displays atlases showing the regional density of each cell type, including total cells, neurons, and PV neurons, along with ratios such as neuron/total cells and PV/total neurons **(E)** The eLemur 3D Atlas tab provides an interactive 3D mesh visualization, allowing users to explore and comprehend the brain’s spatial organization. The interface offers flexible viewing options, an interactive manual, and detailed information about brain regions.

Furthermore, the eLemur platform has been purposefully designed to enhance data reusability, providing researchers with tools to integrate available information into their own projects (see **Methods**). This crucial feature supports ongoing studies in the field by facilitating the incorporation of existing data into new research endeavors.

## Discussion

We present eLemur, a platform of multiplex IHC images and atlases of the mouse lemur brain, enabling detailed exploration of its neuroanatomy at cellular and molecular levels. Our platform provides whole-brain sets of high-resolution immunostained images and 2D/3D reference atlases built through multiplex histology-based delineations of brain structures. It also integrates a 3D cell atlas, providing insights into region-by-region cellular compositions, such as the densities and spatial distributions of non-neuronal, neuronal, and PV-expressing cells across diverse brain regions. eLemur serves not only as a comprehensive platform for advancing the use of the mouse lemur as a model organism in neuroscience but also facilitates multimodal data integration, comparative analysis, and further discoveries.

Compared to previous Nissl-based atlases, the open access to the multiplex IHC images encourages users to further investigate inter-regional expressions of cellular and molecular markers, offering potential findings beyond those explored in this study. For the eLemur atlas, we aimed to achieve high-resolution IHC images and a 3D reference atlas, with 0.65 x 0.65 µm and 10 µm isotropic resolution, respectively. To our knowledge, eLemur provides the richest information on multiple marker proteins with the highest spatial resolution of any available 3D brain atlas of primates, comparable only to the Allen Mouse Brain Atlas^50^.

Additionally, we sought high accuracy in brain section alignment and cell counting. Using block face images collected during whole-brain sectioning, 2D-IHC images could be accurately aligned to restore the original brain section. The accuracy of cell quantification in 3D has been enhanced by calibrating 2D-based cell detection counts with 3D-based cell counts, resulting in more precise density measurements.

Furthermore, through the implementation of deep-learning methods, we showcase computer-assisted region segmentation, which significantly enhances the efficiency of annotating brain structures on microscopy images (**Fig. S4**). In recent years, the rapid development and extensive application of image classification methods using machine learning have revolutionized the analysis of biomedical images, including fluorescent images^58–62^. Despite these advancements, the segmentation of anatomical structures still relies heavily on human expertise to identify the plane of a region through comparison to pre-existing literature, decide the scope of a region through putting together anatomical cues, and assign names to recognized structures. In face of such tedious procedures, our approach offers a significant advancement in segmenting images that share only partial overlapping staining features with those used in the training set through bootstrapping steps. Moreover, we have effectively employed a similar deep learning approach for cell counting and density estimation in individual brain regions, showcasing the versatility of these techniques in quantitative neuroanatomy (**Fig. 4**). While the eLemur atlas represents a significant advancement, several challenges and limitations remain. Tissue distortions, anatomical variations, and technical constraints inherent in brain mapping techniques underscore the necessity for continued refinement and expansion of the atlas. Future research endeavors should prioritize delineating further detailed brain structures, integrating additional markers and imaging modalities, and addressing the heterogeneity of brain structures across individuals. The reliance on a single reference specimen for annotation may inadequately capture the anatomical variability present across diverse individuals and populations of mouse lemurs.

In recent decades, neuroscience research has predominantly focused on a narrow selection of species, often overlooking the vast biological diversity across the animal kingdom^15^. Despite the increasing recognition of the limitations inherent in rodent models, the adoption of alternative model systems has been relatively slow. However, incorporating diversity in animal models is essential for understanding the breadth of biological mechanisms across species. The mouse lemur presents a distinct advantage in this regard. It bridges the gap between the rich array of tools and insights derived from rodent-based studies and the need for models that more closely align with human biology. As a non-human primate, the mouse lemur shares closer evolutionary and neurobiological similarities with humans than rodents do. Integrating the mouse lemur into neuroscience research allows scientists to explore unique evolutionary adaptations and neurobiological characteristics specific to primates, while also benefiting from the practical advantages of working with a small, easily maintainable species.

In conclusion, the mouse lemur emerges as a promising animal model in neuroscience research due to its potential to bridge the gap between existing mouse studies and non-human primates/humans. By embracing the biological diversity offered by the mouse lemur, researchers can explore new frontiers in neuroscience and address the complex questions underlying brain function across species. The eLemur atlas serves as a valuable resource for the neuroscience community, offering detailed insights into the neuroanatomy of the mouse lemur brain.

## Materials and Methods

### Animals

Five mouse lemur brains (from 1 male and 4 females) born and raised in the laboratory colony of UMR 7179 (CNRS/MNHN, Brunoy, France; license approval n° A91.114.1) were used in this study. The animals were maintained in cages enriched with branches and wooden nest boxes at a standard temperature of 24– 26°C and relative humidity of 55%. The animals were fed with fresh fruits and a laboratory-made porridge of cereals, milk and eggs. Water and food were available *ad libitum*. Animals were tested in summer-like photoperiod (14 h of light/day). The age range was 11-45 months (mean ± s.d. 26 ±14.697), which is considered as young to middle-aged adults ^63^. All animal care and experimental procedures were approved by the University of Geneva and French ethics committee “Comité d’éthique Cuvier” (authorization APAFIS#2083-2015090311335786). The delivery of the brain specimens was approved in accordance with the convention on international trade in endangered species of wild fauna and/or flora (CITES) since histological experiments were partially performed in Korea (export permit FR1809100082-E and import permit ES2018-03258).

### Tissue preparation, block face imaging, histology, and microscopy

Mouse lemurs were anesthetized and perfused transcardially with 0.1 M phosphate-buffered saline (PBS) and 4% paraformaldehyde in 0.1M phosphate buffer (PFA). Brains were post-fixed in 4% PFA overnight and incubated in 20% sucrose in PBS at 4 °C for cryoprotection. Brains were sectioned coronally at 50 µm thickness (approximately a total of 360 sections per brain) on a freezing microtome (Fisher Scientific HM450). During sectioning, block face images (cutting planes) of the entire brains were photographed with a CMOS camera (Leica IC90 E, image size of 2592px□×□1944px) mounted on a stereomicroscope (Leica M60).

For immunofluorescence, brain sections at a 100-µm interval were permeabilized in 0.3% Triton X-100 in tris-buffered saline (TBS) and blocked in 3% normal goat serum, 3% bovine serum albumin, and 0.3% Triton X-100 in TBS. The sections were incubated with primary antibodies overnight at 4 °C (See **Table S1** for the details of antibodies used). After washing, sections were incubated with secondary antibodies for 3 h at room temperature and counterstained with DAPI. Sections were mounted with mounting media (Vector Labs, H-1400). Secondary antibodies (1:1000) used were Alexa Fluor 488 goat anti-rabbit IgG (Invitrogen, A11034), Alexa Fluor 488 goat anti-mouse IgG (Invitrogen, A11029), Alexa Fluor 555 goat anti-mouse IgG (Invitrogen, A21424), Alexa Fluor 555 goat anti-rabbit IgG (Invitrogen, A21428), Alexa Fluor 633 goat anti-guinea pig IgG (Invitrogen, A21105), and Alexa Fluor 633 goat anti-rat IgG (Invitrogen A21094). Mice were anesthetized, and brains were perfused using the same procedures as above. Mouse brains were post-fixed in 4% PFA overnight and sectioned coronally at 50 µm thickness (approximately 240 sections per brain) on a vibratome (Leica VT1200S). Brain sections containing regions of interest were permeabilized in 1% Triton X-100 in TBS and blocked in 5% normal goat serum, goat anti-mouse Fab fragments (Jackson Lab, 715-007-003) and 0.4% Triton X-100 in TBS. Identical antibodies and procedures for immunohistochemistry were used as above. Widefield images were acquired using an Axioscan Z1 slide scanner (Carl Zeiss Microscopy) equipped with a 10X 0.45 NA Plan-Apochromat air lens. For cell counting analysis, confocal images were obtained at 0.54 μm depth intervals using the LSM 780 confocal microscope (Carl Zeiss Microscopy) equipped with a 40x 1.4 NA Plan Apochromat oil lens.

### Shading correction

A custom python script was used to apply the BaSiC algorithm^64^ for flat-field estimation and shading correction on 2D-histology images. To ensure even illumination across sections within a single brain image set, we randomly extracted 20 image tiles (2040px × 2040px) from each section, combining them to estimate flat-field profile using BaSiC with default parameters. The estimated flat-field profile was then resized to match the original image size. A single flat-field profile was then used to correct shading in all image sections independently.

### 2D Block face Alignment

Using a custom-built MATLAB GUI, 2D histology image series were manually inspected for missing or damaged tissues and mispositioning, including flipping and rotation. Rotated or flipped images were preprocessed to center them in the correct orientation, providing better initialization for the following registration process. Tissue areas in both histology and block face images were semi-automatically labeled, with adaptive thresholding followed by manual correction. For images containing multiple disjoint tissues, correspondence between histology and block face images were determined manually by human experts.

Tissue part in the histology images were aligned to the corresponding parts in the block face image via multi-scale 2D rigid registration from Advanced Normalization Tools^65^ in Python.

### Semi-automatic & manual annotation

A custom software with a graphical user interface (GUI) was developed to enable manual annotation of brain regions through cubic spline curves, defined by control points. Annotations were made using a computer mouse or an Apple Pencil as the input device (**Fig. S4C**). Cubic spline curves, due to their scale-invariant nature, allowed for efficient storage and flexible editing of large-volume annotations. 13 major brain regions were manually annotated, and this data was utilized to train a deep learning segmentation network to automate the annotation of unlabeled images (**Fig. S4D**). Due to the complex morphology of brain regions, a PointRend model^66^ with a ResNeXt-101-32x8d backbone was employed. This model was trained for 192,000 iterations on 4 NVIDIA Tesla V100 GPUs. A learning rate initially set at 0.01 was reduced by a factor of 10 at the 48,000th, 128,000th, and 144,000th iterations. Data augmentation methods such as vertical and horizontal flips, random scaling, rotation, and channel shifts were applied. After fully annotating the major regions throughout the dataset, subregions were delineated by drawing virtual ’cut lines’ on the annotated major brain regions (**Fig. S4E**). The ’cut lines’ were utilized to segment the major brain regions, and each resulting section was manually assigned the appropriate region ID, resulting in the finalized brain region annotations.

### 3D Template atlas generation

To generate initial 3D template atlas volume, additional 2D non-linear multi-scale registration was performed between the binary mask images of histology and corresponding block face image, and later applied to the 2D label images. Symmetric diffeomorphic registration^67^ was used to interpolate 10 sections between two consecutive label images, achieving a 10 µm resolution along anterior-posterior direction.

Interpolation was applied independently to each brain region. Since both initial non-linear registration and interpolation were incomplete, interpolated regions had overlaps. In such cases, overlapping voxels were unassigned from either region, and each region was equally expanded iteratively until the boundaries touched and filled the gap. Once the 3D isotropic (10 x 10 x 10µm) label volume with no overlapping was ready, a 3D mesh for each region was generated using the marching-cube algorithm^53^. To generate a smoother isosurface, a Laplacian filter (λ =1e-4, iteration=100) was applied to the 3D mesh of each region.

Consequential region overlapping was resolved by transforming the smoothed mesh back to the 3D volume image and overlapped voxels were fixed within voxel space. The smoothing parameters of Laplacian filters were gradually decreased each step of iterations to maintain boundary sharpness.

### MRI

A publicly available dataset of a population averaged T2-weighted MR brain template^32^ was used to annotate 19 Brodmann areas. The population averaged T2-weighted MR brain template and corresponding annotation labels were registered onto 3D block face volume via multi-level registration consisting of a series of 3D affine and non-linear transformation using Advanced Normalization Tools^65^ in Python.

Cortical regions from a 2D slice of registered annotation labels were registered onto cortical areas in the corresponding eLemur slice via 2D deformable registration. The transferred annotations were manually refined to ensure the boundaries between regions lay parallel to radial direction.

### Cell detection model training and validation

The Detectron2 framework^68^ was employed to train the model for DAPI and NeuN detection. The training dataset comprised 20 images and 3646 cell annotations. A Mask R-CNN model with a ResNeXt-101-32x8d backbone, initialized with the weights of the model pretrained on the MS-COCO dataset, was utilized. This model was trained for 6000 iterations on two NVIDIA Tesla V100 graphics processing units (GPUs) with a batch size of 6. A fixed learning rate of 0.01, along with weight decay and gradient norm clipping, was applied. Vertical and horizontal flipping, and random scaling were applied for image augmentation. For PV cell detection, which exhibit a distinct appearance from DAPI and NeuN, a separate model with the same architecture was trained on a dataset containing 77 images with 775 annotations, using the same hyperparameters and procedures as the DAPI and NeuN model, but with an adjusted batch size of 8. To benchmark our model against Cellpose 2.0, we followed the hyper-parameters described in the original paper^56^ to fine-tune specialist Cellpose models, utilizing our two distinct training datasets for DAPI/NeuN and PV cell types. The fine-tuning process was conducted through the Cellpose 2.0 training API, with a learning rate of 0.1, for a total of 500 epochs. All other parameters remained at their default configurations. This training utilized a single Tesla V100 GPU equipped with 32GB of memory.

For the validation of cell detection models, image patches of 166.4 x 166.4μm were randomly selected from various brain regions and annotated based on consensus from six experts. The ‘cellpose.metrics.average_precision’ function from the Cellpose 2.0 evaluation API was used to calculate average precision, recall rate, and the F1-score. Masks with an Intersection over Union (IoU) of 0.5 or greater compared to ground-truth labels were considered true positives, while those below this threshold were labeled as false positives. Non-detected ground truths were classified as false negatives. A total of 29 annotated image patches were used for validating the models for DAPI and NeuN detection. The Cellpose 2.0 model’s performance was further assessed by training with 18 additional image patches, with the remaining 11 patches serving as the test set, demonstrating performance enhancement with more training data. For PV detection, 9 image patches were used to assess the performance of both our model and the Cellpose 2.0 model.

### eLemur website development and data access

The eLemur website has been developed to provide neuroscientists with necessary tools for integrating high-resolution mouse lemur brain atlas and data into their own research. Customized Vue.js components were developed to construct the platform’s user interface, while the Neuroglancer^73^ engine powers the full resolution imaging pages, offering detailed navigation. These components are open-source and accessible at our GitHub repository (https://github.com/feng-lab/zjbrainscience-front/tree/eeum), promoting transparency and collaboration. The 3D atlas of the mouse lemur brain is available through the eLemur website and consists of 3 files:

1. eLemur_3d_atlas_mesh.zip: 3D meshes of the anatomical structures
2. eLemur_3d_atlas.mhd.zip: volumetric atlas files
3. eLemur_3d_atlas.label: label file specifying the ID, color code and name of each anatomical structure Access to the eLemur image datasets is facilitated through the Tensorstore^69^ library, which allows for downloading the whole dataset or a subset of the data. An example of how the Tensorstore library can be used to access and download a subset of the image data as well as its brain region annotation is provided below:

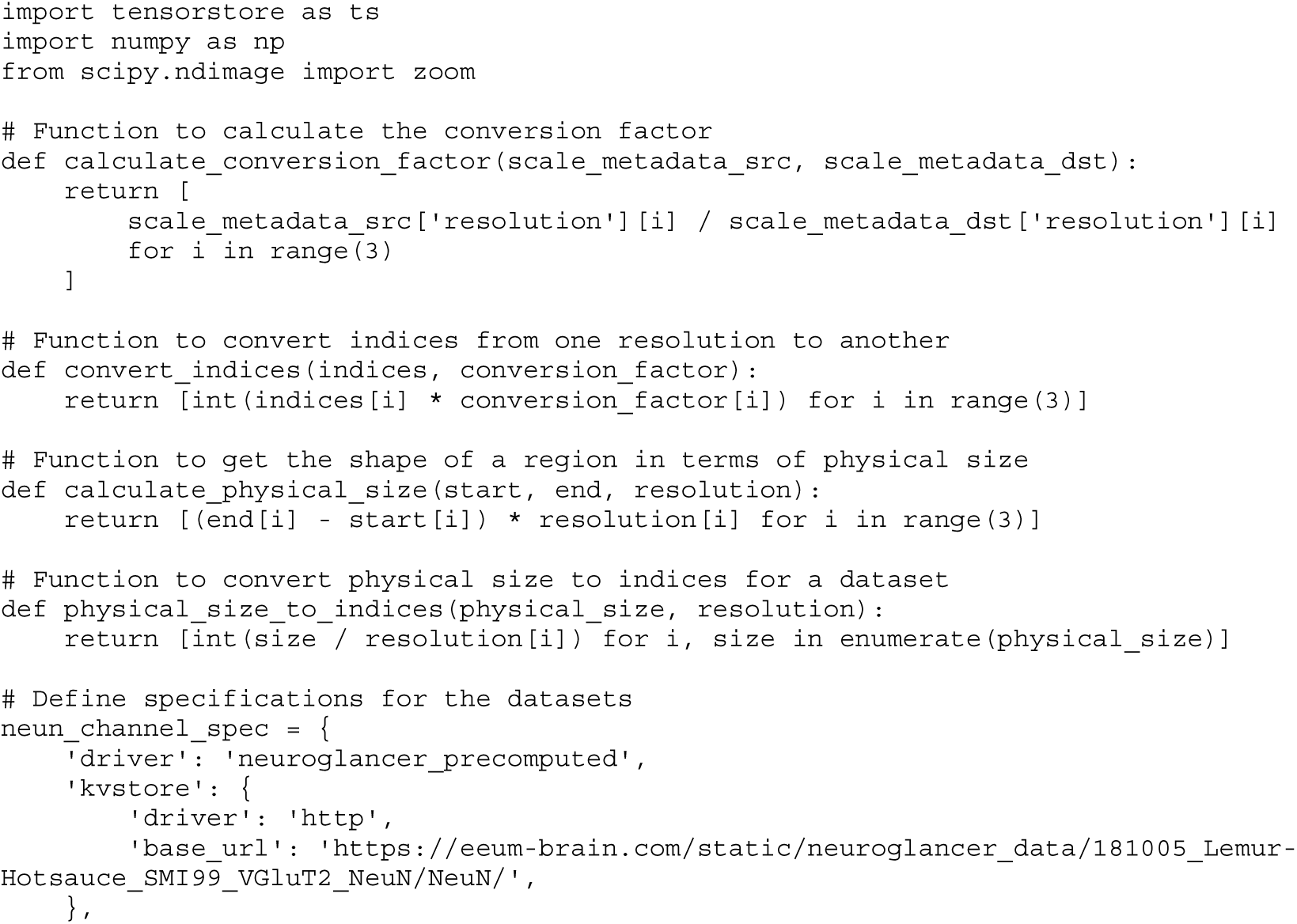
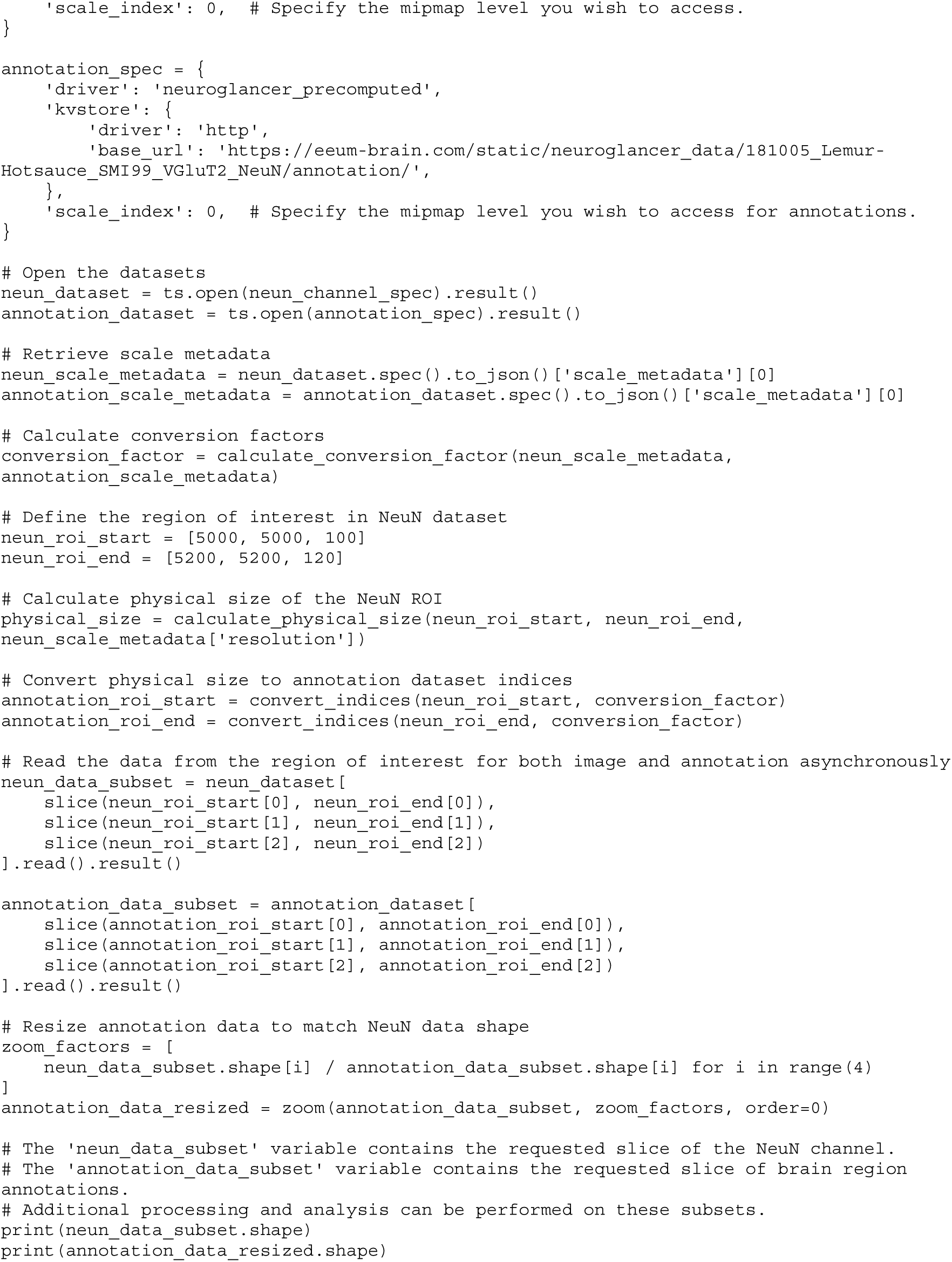

## Acknowledgements

This work was supported by the Human Frontiers Science Program (grant no. RGP0024/2016) and the KIST intramural program (2E32901). We thank Ziao Liu for training and benchmarking cell detection models, Qin Zhu, Qi Tian, Xinhang Li, Cong Wang for assistance with cell detection validation, Wen Wu for developing the online interface for eLemur, and Sohyeon Jeon for contributions to brain region annotation.

## Author Contributions

Jinhyun K. and D.H. designed research; Jiwon K., A.C.L.H., and Jayoung K. performed research; Jiwon K., Jayoung K., H.J., and L.F. analyzed data; F. P. contributed to animal care and use; H.J., L.F., and Jinhyun Kim developed website-based resources; and Jayoung K. and Jinhyun K. wrote the paper with contributions from H.J., Jiwon K., and L.F, and feedback from all authors.

## Competing Interest Statement

The authors declare no competing interest

---

**Supplementary Figure 1.**
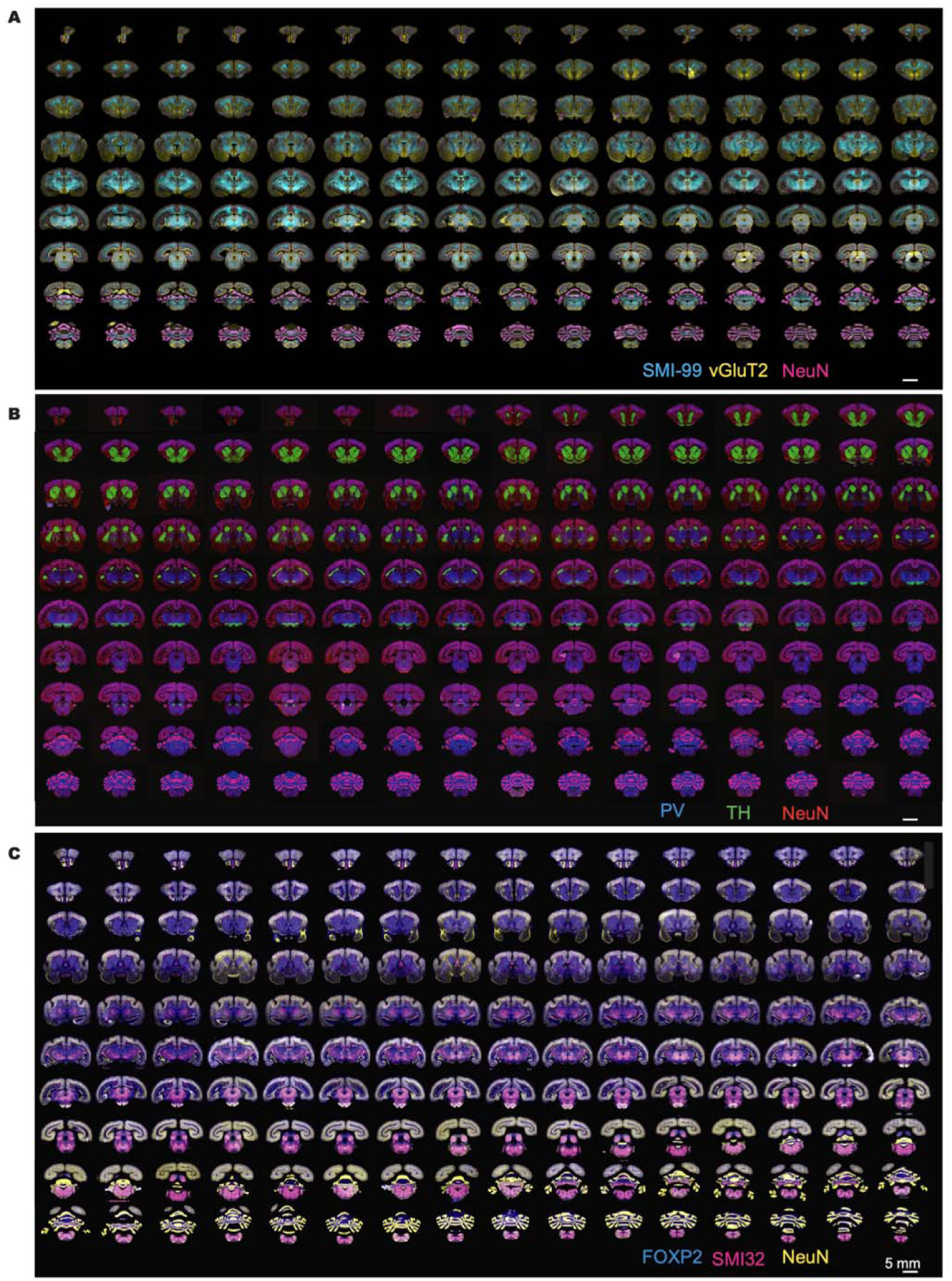
Whole-brain immunofluorescence images of the mouse lemur. Gallery view of whole-brain IHC sets of the mouse lemur brain stained with **(A)** SMI-99, VGLUT2, NeuN, **(B)** PV, TH, NeuN, and **(C)** FOXP2, SMI-32, and NeuN.

**Supplementary Figure 2.**
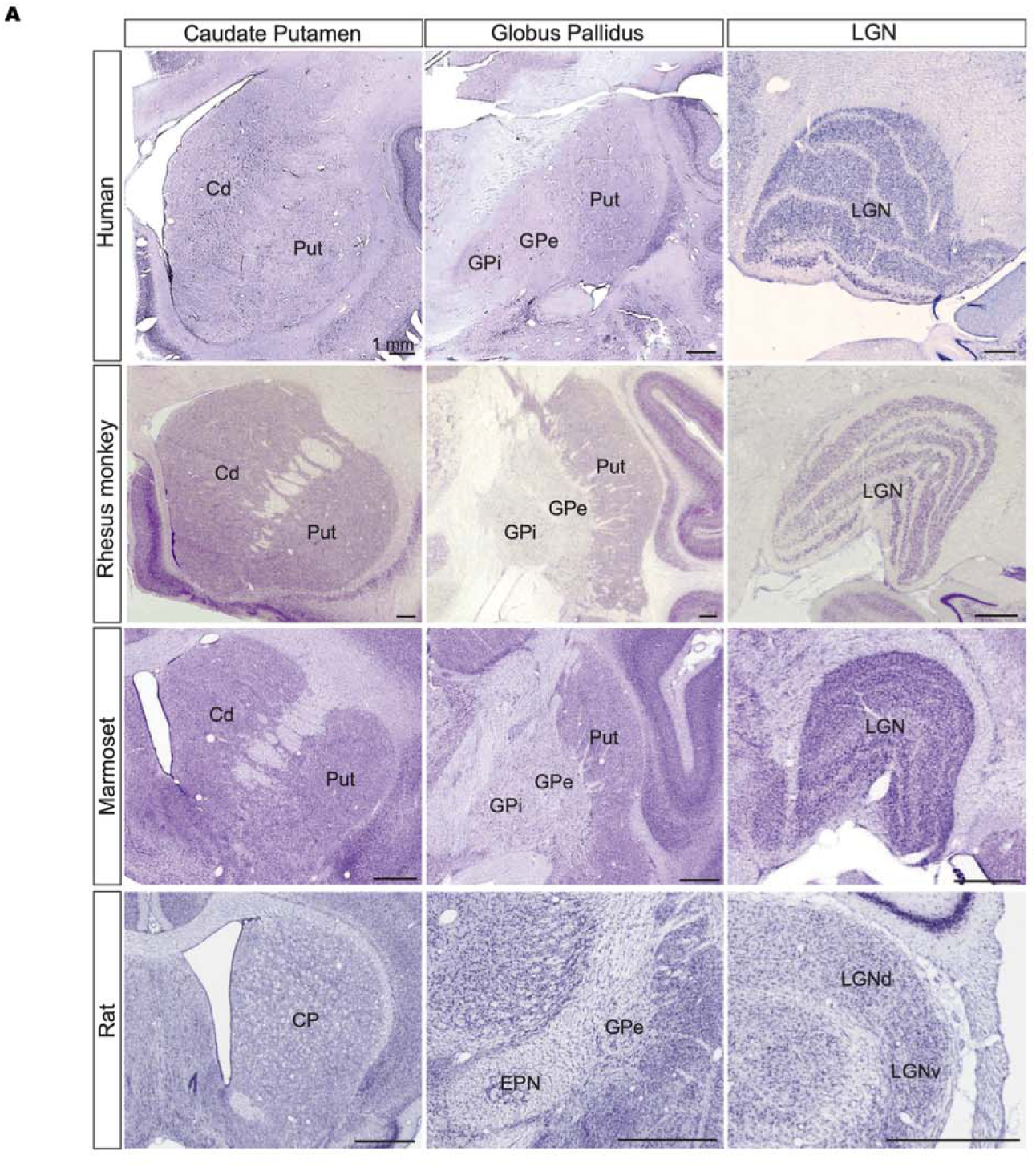
Comparative analysis of the basal ganglia and LGN across species. **(A)** Comparison of the basal ganglia and LGN across species utilizing publicly available Nissl staining-based atlases. Human data were extracted from the Allen Reference Atlas, while stainings for rhesus monkey, marmoset, and rat were sourced from Brainmaps.org. In primates, the basal ganglia exhibit distinct caudate and putamen segments separated by a fiber track, alongside a layered structure of the LGN. Conversely, in rodents, organization of the basal ganglia and LGN in rats differs from that in primates (also see Figure 1D).

**Supplementary Figure 3.**
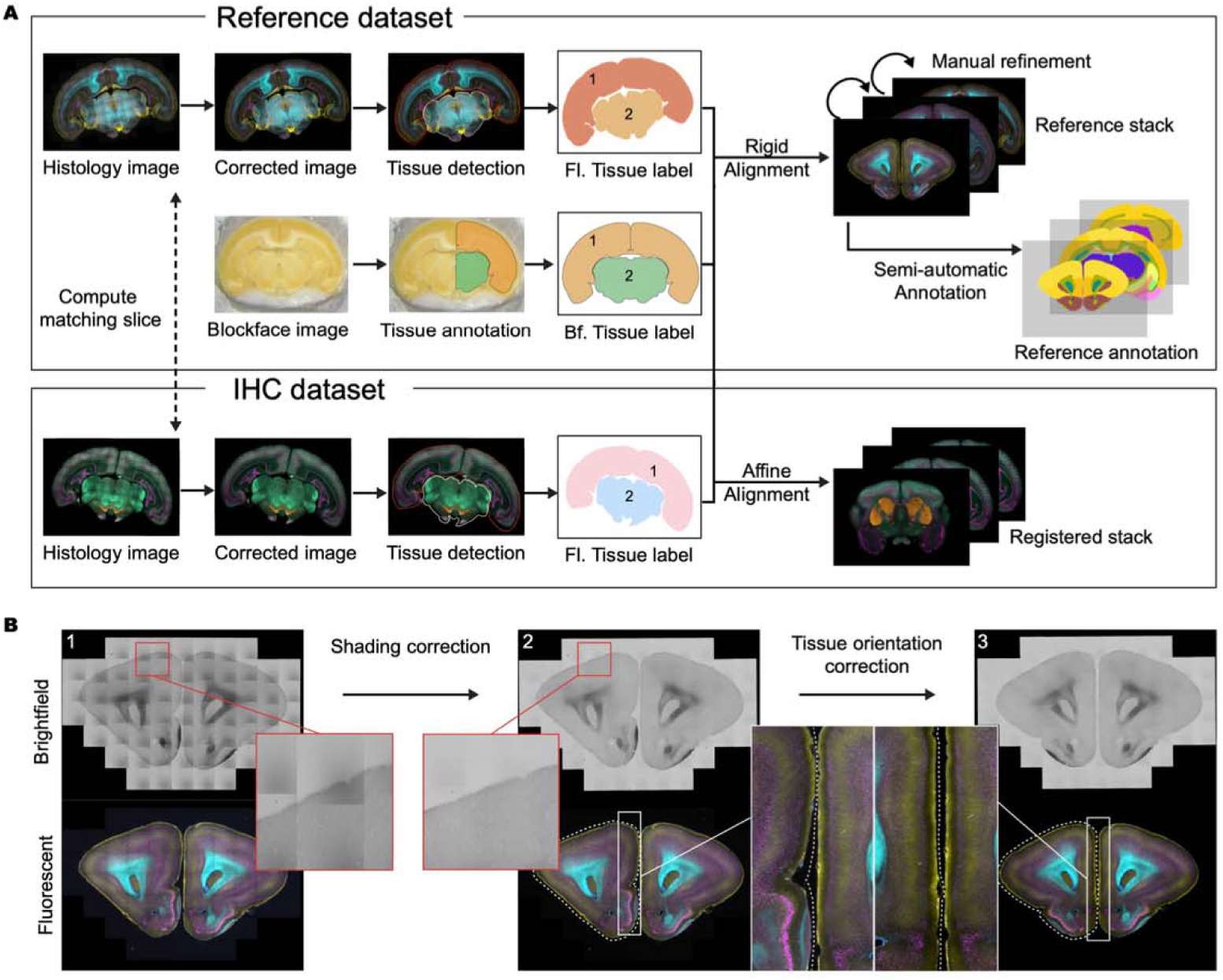
Image processing pipeline for the immunofluorescent image dataset. **(A)** The flowchart outlines the steps involved in processing and aligning the reference and other immunohistochemistry (IHC) datasets. Immunofluorescent images first undergo correction and alignment to the corresponding block face image. Subsequently, semi-automatic region annotation is conducted. To minimize artifacts arising from deformation, only rigid transformations (rotation and translation) are applied to the reference dataset. **(B)** A coronal view of a brain slice showing three stages of image processing: The original image before any correction procedures (left), the image after shading correction to enhance clarity and remove irregular illumination (middle), and the final image after undergoing repair procedures to address any remaining artifacts or imperfections (right).

**Supplementary Figure 4.**
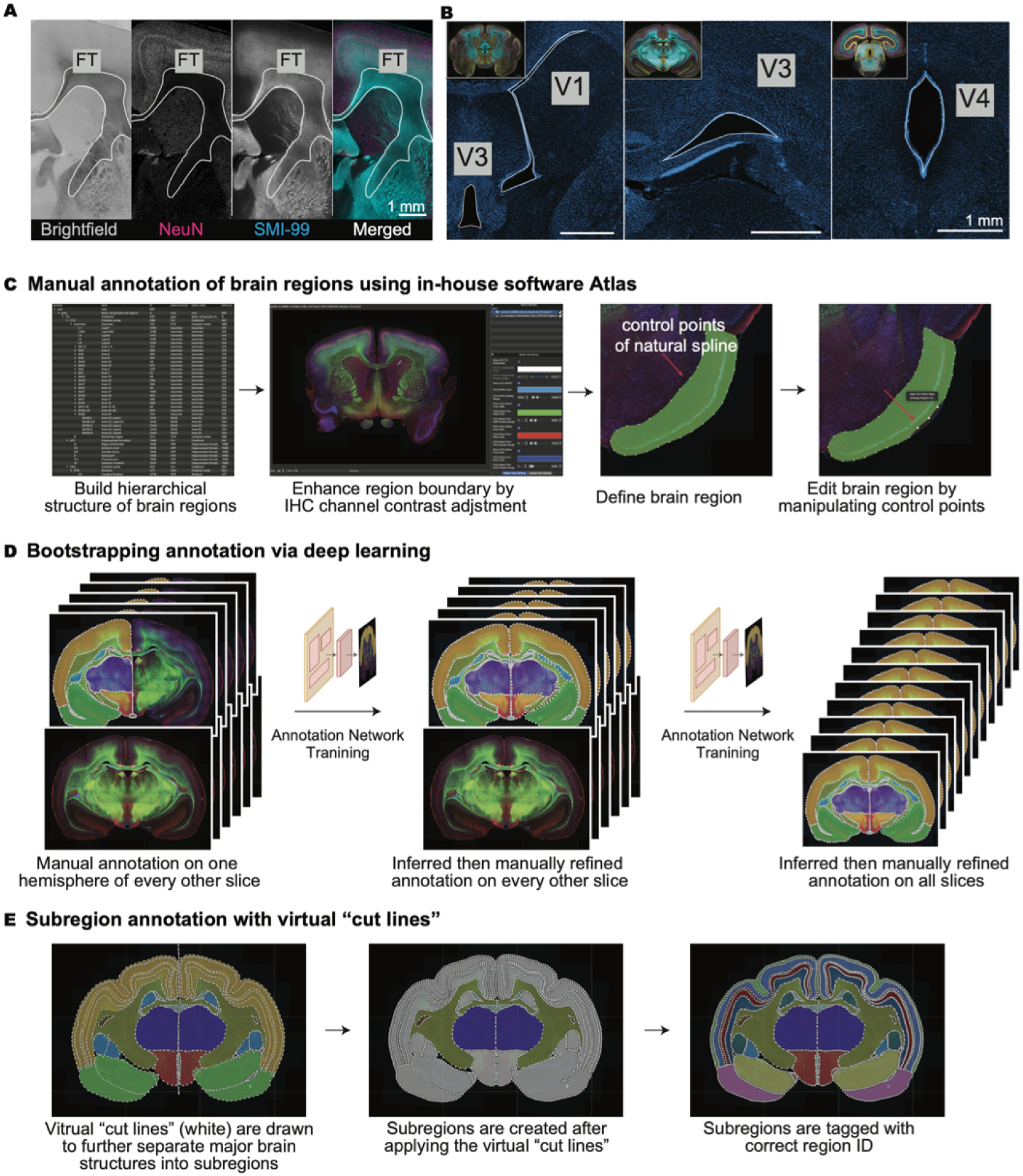
Annotation of brain structures. **(A, B)** Fiber tracks and ventricles are delineated using multiplex immunosignals, complemented by brightfield contrast images. **(C)** Manual annotation of brain regions was performed using an in-house annotation software featuring a Graphical User Interface (GUI). Brain regions were delineated with closed natural cubic splines. **(D)** Annotation bootstrapping through deep learning involved training two deep region segmentation networks to propagate partial annotations across the entire stack. **(E)** Subregion annotation was achieved using virtual "cut lines" to define smaller subregions by separating major brain regions.

**Supplementary Figure 5.**
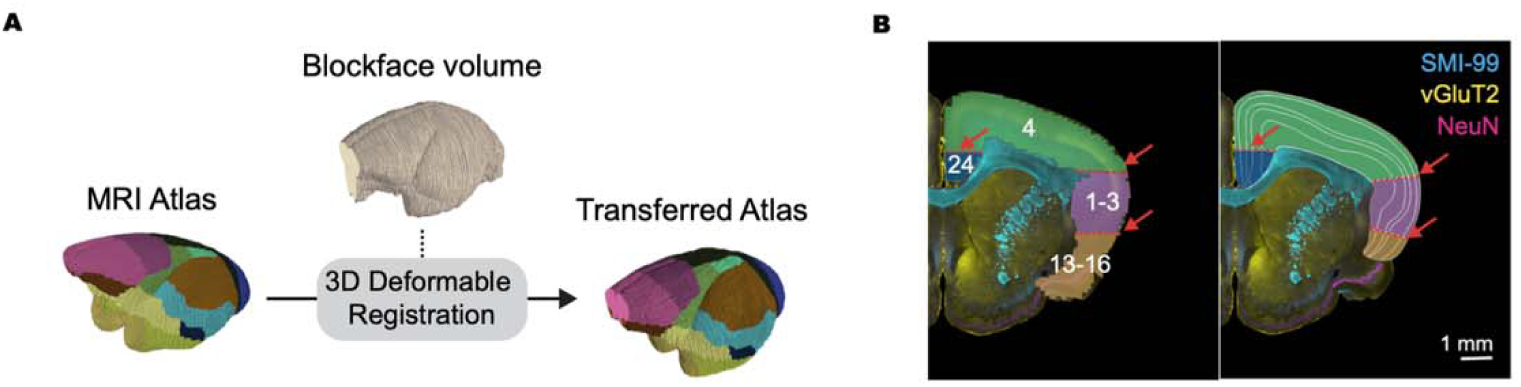
Integration of the MRI-based atlas into 2D immunofluorescence images. **(A)** Flowchart of the population averaged T2-weighted MRI atlas registration to the block face volumetric image. **(B)** Slice-by-Slice manual correction and refinement of registered MRI atlas. Deformed atlas is overlaid on to corresponding IHC slices and the boundaries between Brodmann area (red) are refined to 1) match cellular architectures and 2) be aligned parallel to the radial direction.

**Supplementary Figure 6.**
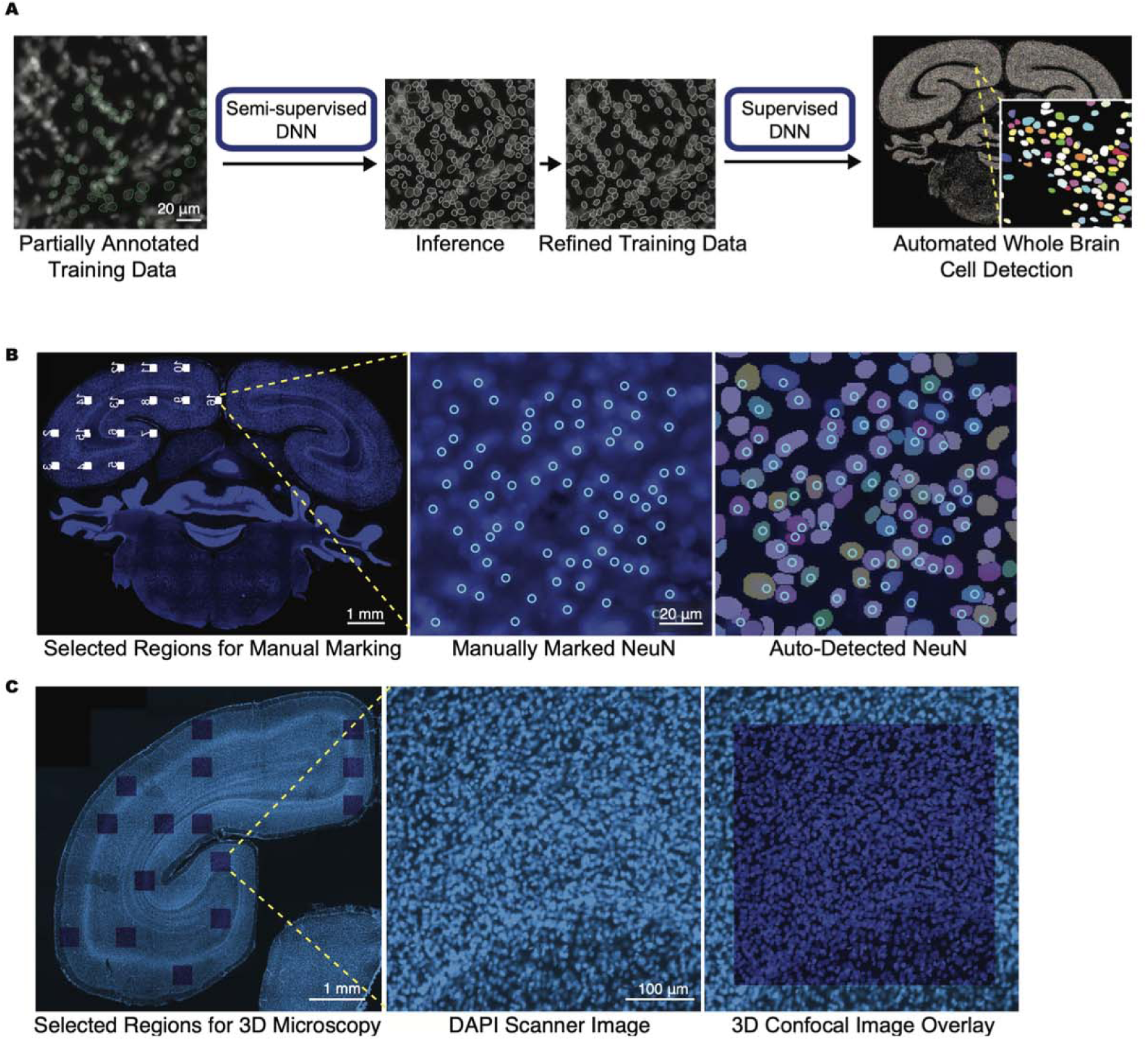
Automated whole-brain cell detection algorithm. **(A)** Workflow of automated whole-brain cell detection algorithm based on partially annotated data. **(B)** Validation of automatic NeuN-positive neuron detection results. Small image patch is sampled uniformly within the slice (left). For each patch, NeuN cells are manually annotated by human experts (center), and automatic detection results are validated against manual detections (right). **(C)** Validation of automatic 2D DAPI-positive cell detection results against 3D confocal cell counts. Small image column is sampled from 3D confocal image and corresponding image patch from 2D IHC slice is determined (left). Detection results from 2D slice and compared and validated with 3D confocal image (center and right).

**Supplementary Figure 7.**
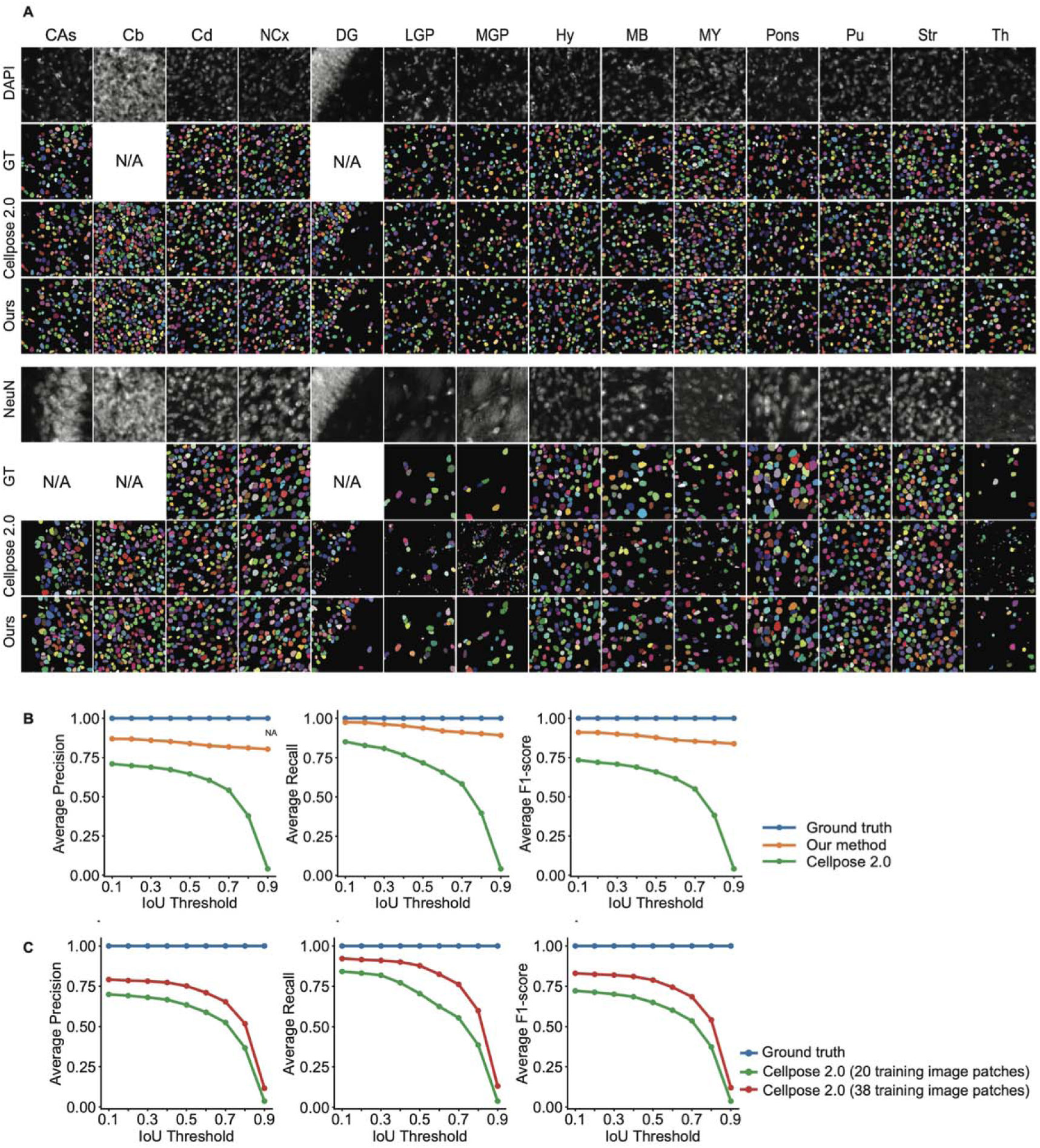
Validation of DAPI and NeuN detection algorithm. **(A)** Random selected image patches from various brain regions, showing consensus annotations from six experts as ground-truth (GT)”, detection results from Cellpose 2.0, and our method. **(B)** Average precision, recall, and F1-score for different cell detection methods, demonstrating that our method outperforms Cellpose 2.0. **(C)** Performance enhancement of Cellpose 2.0 with more training datasets.

**Supplementary Figure 8.**
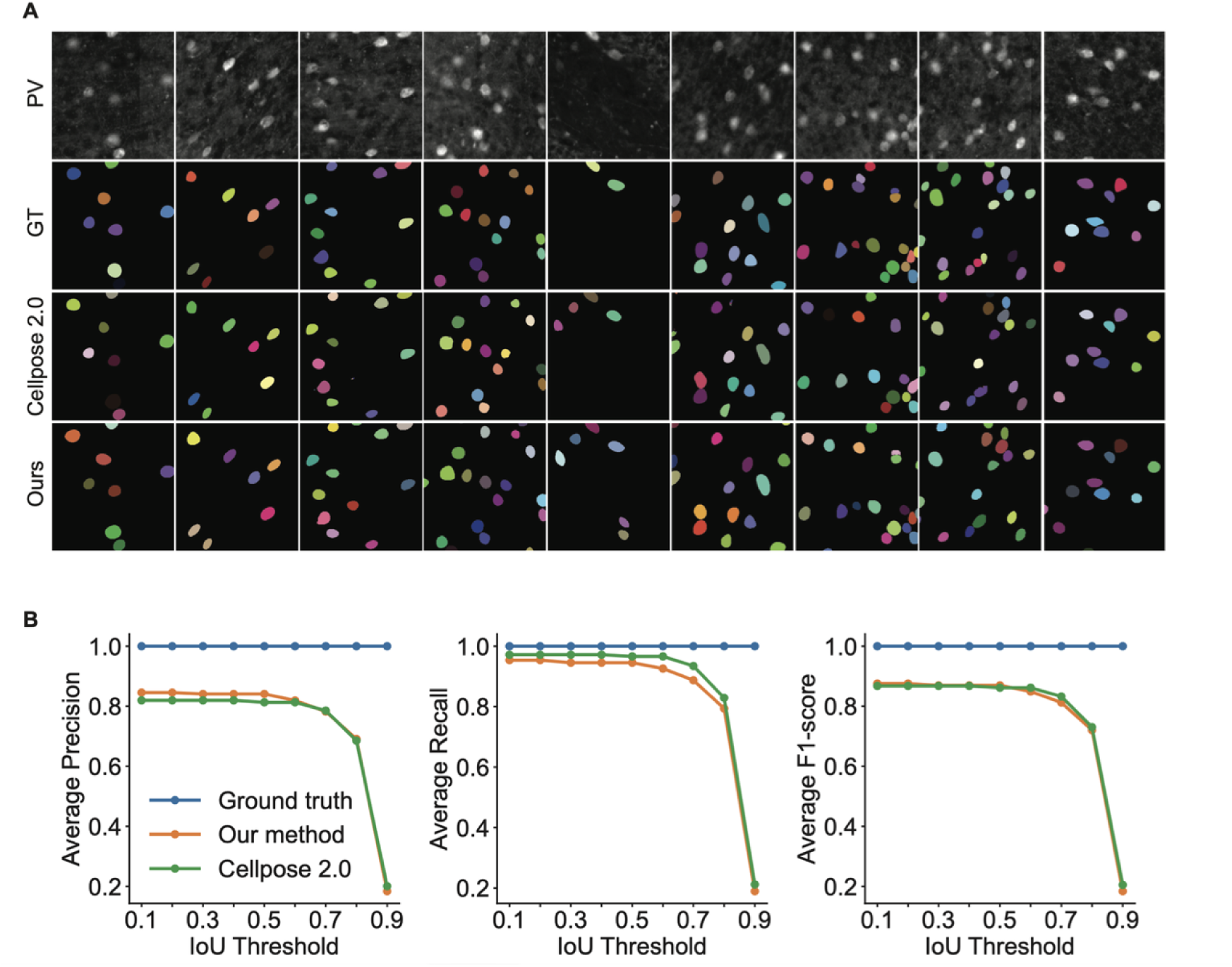
Validation of PV detection algorithm. **(A)** Random selected image patches from the PV channel, showing manual annotations, detection results from Cellpose 2.0, and our method. **(B)** Average precision, recall, and F1-score for different cell detection methods.

**Table S1.** Primary antibody information.

| Stain | Target | Host | Vendor | Cat. No. | RRID | Dilution |
| --- | --- | --- | --- | --- | --- | --- |
| NeuN | Neuronal nuclear antigen | guinea pig | Millipore | ABN90P | AB_2341095 | 1:1000 |
| FOXP2 | Forkhead box protein P2 | rabbit | Abcam | AB16046 | AB_2107107 | 1:1000 |
| PV | Parvalbumin | mouse | Swant | 235 | AB_10000343 | 1:1000 |
| TH | Tyrosine hydroxylase | rabbit | Millipore | AB152 | AB_390204 | 1:1000 |
| SMI-99 | Myelin basic protein | mouse | BioLegend | 808401 | AB_2564741 | 1:1000 |
| SMI-32 | Neurofilament H | mouse | BioLegend | 801702 | AB_2715852 | 1:1000 |
| vGluT2 | Vesicular glutamate transporter 2 | rabbit | Synaptic Systems | 135403 | AB_887883 | 1:1000 |

Information on the primary antibodies used to generate mouse lemur whoa-brain immunohistochemistry (IHC) sets. The antibodies were selected based on their compatibility with mouse lemur brain tissues after initial testing on mouse brain tissues.

**Table S2.**
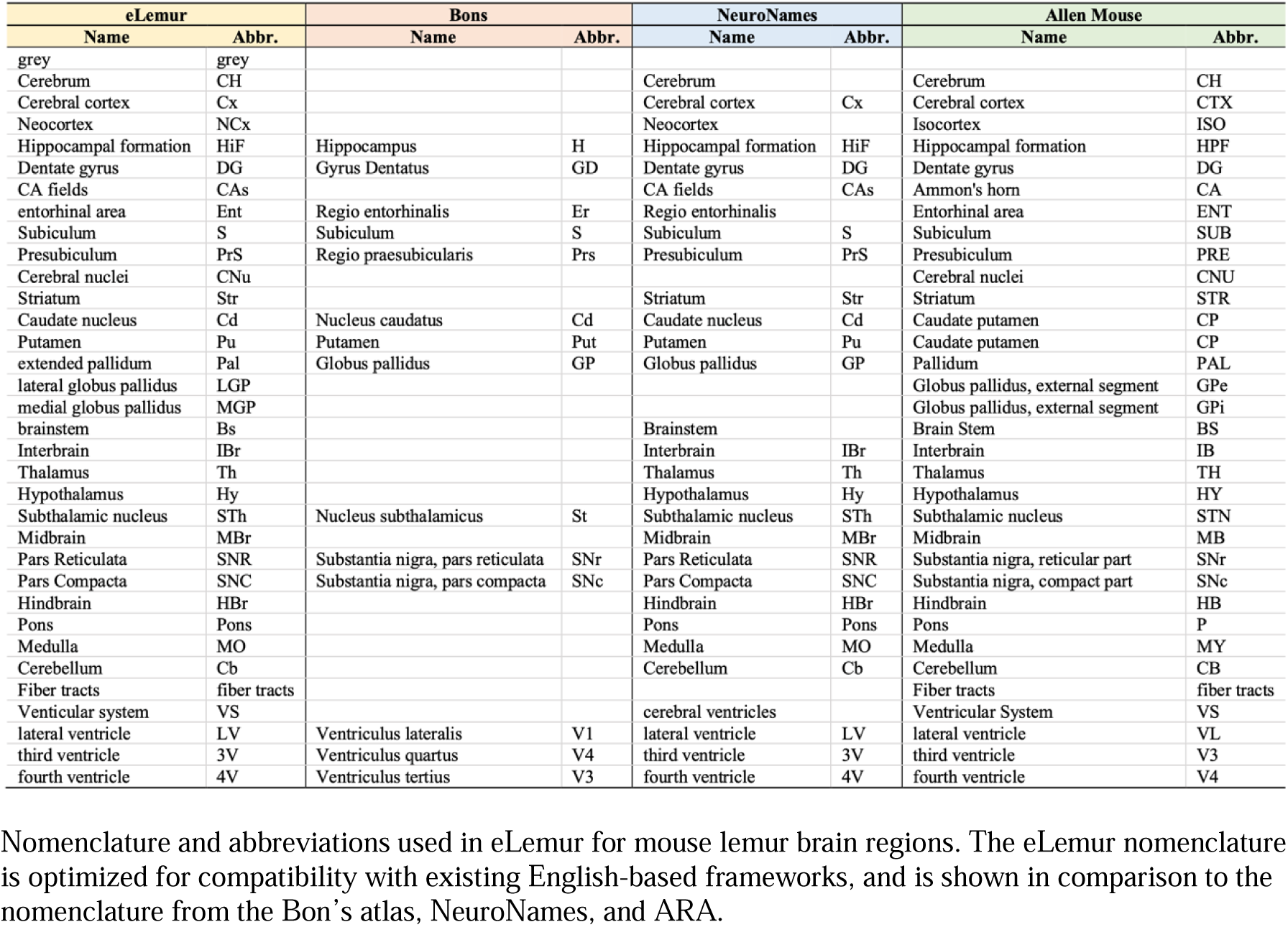
Mouse lemur brain region nomenclature.

**Table S3.**
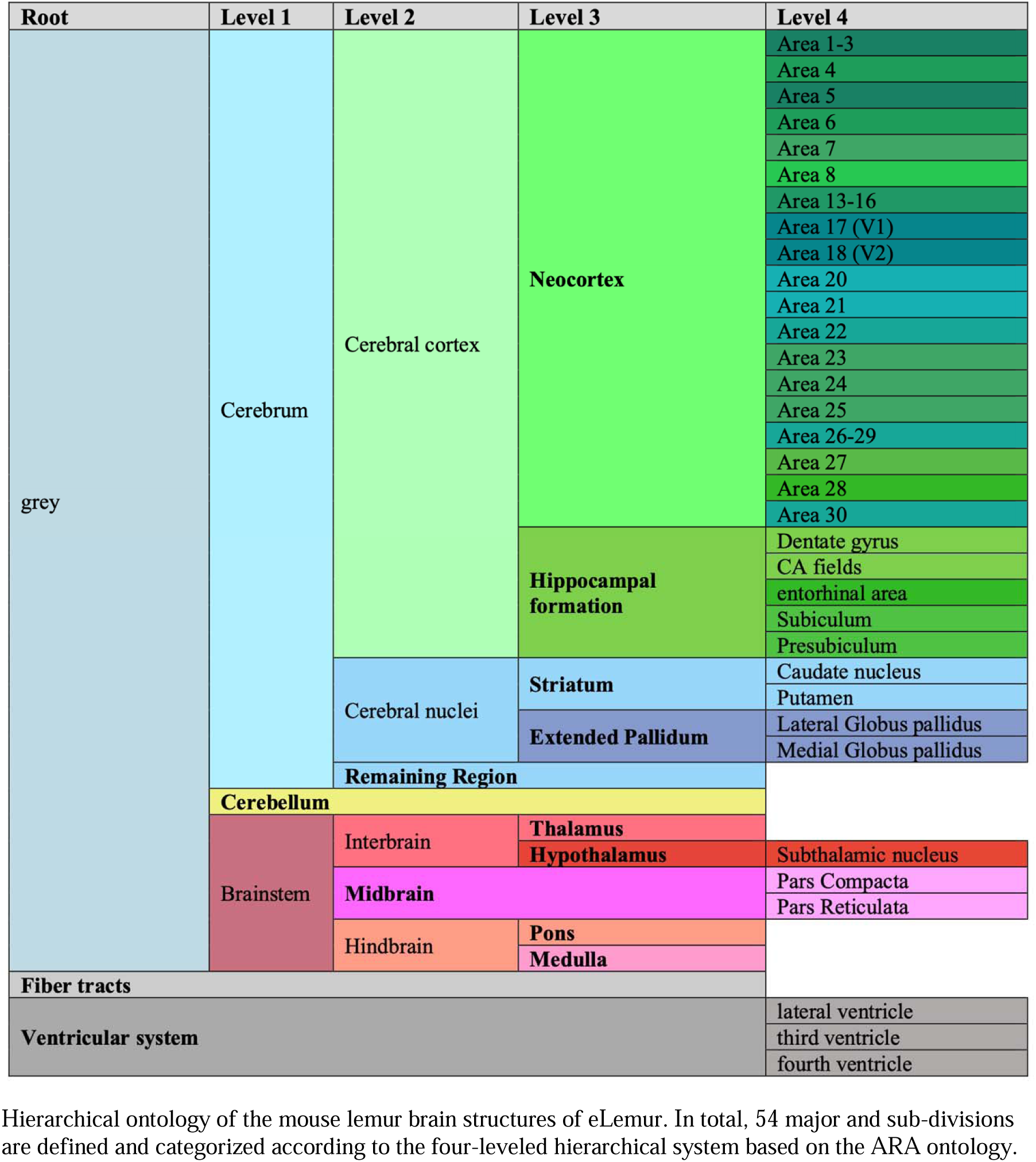
Mouse lemur brain ontology.

## Notes

https://eeum-brain.com/#/lemurdatasets

### Competing Interest Statement

The authors have declared no competing interest.

